# Tirant stealthily invaded natural *Drosophila melanogaster* populations during the last century

**DOI:** 10.1101/2020.06.10.144378

**Authors:** Florian Schwarz, Filip Wierzbicki, Kirsten-André Senti, Robert Kofler

**Author notes:** to whom correspondence should be sent. Institute of Molecular Biotechnology of the Austrian Academy of Sciences (IMBA), Vienna Biocenter (VBC), Dr. Bohrgasse 3, 1030 Vienna, Austria.

## Abstract

It was long thought that solely three different transposable elements - the I-element, the P-element and hobo - invaded natural *D. melanogaster* populations within the last century. By sequencing the ‘living fossils’ of *Drosophila* research, i.e. *D. melanogaster* strains sampled from natural populations at different time points, we show that a fourth TE, Tirant, invaded *D. melanogaster* populations during the past century. Tirant likely spread in *D. melanogaster* populations around 1938, followed by the I-element, hobo, and, lastly, the P-element. In addition to the recent insertions of the canonical Tirant, *D. melanogaster* strains harbour degraded Tirant sequences in the heterochromatin which are likely due to an ancient invasion, possibly predating the split of *D. melanogaster* and *D. simulans*. In contrast to the I-element, P-element and hobo, we did not find that Tirant induces any hybrid dysgenesis symptoms. This absence of apparent phenotypic effects may explain the late discovery of the Tirant invasion. Recent Tirant insertions were found in all investigated natural populations. Populations from Tasmania carry distinct Tirant sequences, likely due to a founder effect. By investigating the TE composition of natural populations and strains sampled at different time points, insertion site polymorphisms, piRNAs and phenotypic effects, we provide a comprehensive study of a natural TE invasion.

## Introduction

Transposable elements (TEs) are DNA sequences that multiply within host genomes, even if this activity is deleterious to hosts (Doolittle and Sapienza, 1980; Orgel and Crick, 1980; Hickey, 1982; Wicker et al., 2007). To enhance their rate of transmission into the next generation, TEs need to infect the germ cells. While most TEs achieve this by being active in the germline, some LTR retrotransposons generate virus-like-particles in the soma surrounding the germline, which may infect the germ cells (Song et al., 1997; Blumenstiel, 2011; Goodier, 2016; Moon et al., 2018; Wang et al., 2018). Since many TE insertions are deleterious, host organisms evolved elaborate defense mechanisms against TEs (Brennecke et al., 2007; Marí-Ordóñez et al., 2013; Yang et al., 2017). In *Drosophila melanogaster* the defense against TEs is based on piRNAs, i.e., small RNAs with a size between 23-29nt, that repress TE activity at the transcriptional and the post-transcriptional level (Gunawardane et al., 2007; Brennecke et al., 2007; Sienski et al., 2012; Le Thomas et al., 2013). One option to escape the host defense is to infect a novel species. Many TEs cross species boundaries, e.g. due to horizontal transfer (HT) from one host species to another, and trigger invasions in naive species not having the TE (Mizrokhi and Mazo, 1990; Maruyama and Hartl, 1991; Lohe et al., 1995; Terzian et al., 2000; Sánchez-Gracia et al., 2005; Loreto et al., 2008; Kofler et al., 2015a; Peccoud et al., 2017). A striking example for a high frequency of TE invasions can be seen in *D. melanogaster*, which was invaded by at least three different TE families within the last century: the I-element, hobo and the P-element (Kidwell, 1983; Anxolabéhère et al., 1988; Periquet et al., 1989; Daniels et al., 1990a,b; Bucheton et al., 1992; Bonnivard et al., 2000). All of these three TEs actively replicate only in the germline and induce some phenotypic effects, the hybrid dysgenesis (HD) symptoms, which historically led to the discovery of the recent TE invasions in *D. melanogaster* (Bingham et al., 1982; Biémont, 2010). An important hallmark of these HD symptoms is that the direction of crosses between two strains is important. The offspring of crosses between males carrying a genomic factor (the TE) and females not carrying this factor frequently show various symptoms - such as atrophic ovaries (P-element, hobo), reduced hatching rates (I-element), or male recombination (P-element, hobo) - whereas the offspring of the reciprocal crosses are usually free of symptoms (Kidwell et al., 1977; Bucheton et al., 1976; Blackman et al., 1987; Yannopoulos et al., 1987). Hybrid dysgenesis thus has a cytoplasmic as well as a genomic component. While TEs were quickly identified as the responsible genomic factor, the cytoplasmic component which is solely inherited from the mothers, the piRNAs, was discovered much later (Bingham et al., 1982; Brennecke et al., 2008). It was realized that the presence of a HD-inducing TE in a strain mostly depends on the sampling date of a strain, where more recently sampled strains frequently carry the TE while old strains, sampled before the invasion, do not. It was thus suggested that the HD-inducing TEs recently invaded *D. melanogaster* populations (Kidwell, 1983; Periquet et al., 1994). These invasions were probably triggered by HT events, where the P-element was likely acquired from *D. willistoni* and the I-element as well as hobo possibly from *D. simulans* (or another species from the simulans clade) (Daniels et al., 1990a,b; Simmons, 1992; Loreto et al., 2008). However, even the old strains carried short and highly degraded (probably inactive) fragments of the I-element and hobo, mostly in the heterochromatin (Bucheton et al., 1984, 1986; Daniels et al., 1990a; Bucheton et al., 1992). Hence, the I-element and hobo likely invaded *D. melanogaster* populations at least twice. Solely the P-element does not have any similarity to sequences in the *D. melanogaster* genome, which suggests that the P-element invaded *D. melanogaster* populations for the first time. The *D. melanogaster* strains sampled at different time points, previously labeled as the ‘living fossils’ of *Drosophila* research (Bucheton et al., 1992), were not only used to discover the three recent TE invasions, but also to estimate the timing of the invasions: the I-element invasion occurred presumably between 1930 and 1950, the hobo invasion around 1955 and the P-element invasion between 1950 and 1980 (Kidwell, 1983; Anxolabéhère et al., 1988; Periquet et al., 1989). By sequencing these ‘living fossils’ we discovered an additional, hitherto undetected TE invasion. We show that Tirant invaded natural *D. melanogaster* populations, likely between 1930-1950. Tirant is an LTR retrotransposon and a member of the *Ty3/Gypsy* superfamily (Moltó et al., 1996; Viggiano et al., 1997; Cañizares et al., 2000; Terzian et al., 2001). It encodes an envelope protein and completes the retroviral cycle in the closely related *D. simulans* (Marsano et al., 2000; Lemeunier et al., 1976; Akkouche et al., 2012). *Tirant may thus replicate as an infective retrovirus*.

## Results

Given this striking accumulation of TE invasions within the last century (Kidwell, 1983; Anxolabéhère et al., 1988; Periquet et al., 1989; Daniels et al., 1990a,b; Bucheton et al., 1992; Bonnivard et al., 2000), we speculated that additional, hitherto undetected TEs, may have recently invaded *D. melanogaster* populations. To test this hypothesis, we compared the abundance of TEs between one of the oldest available *D. melanogaster* laboratory strains, Canton-S (collected by C. Bridges in 1935, Lindsley and Grell (1968)) and the reference strain, Iso-1 (fig. 1A; (Brizuela et al., 1994)). We aligned publicly available short read data from these strains to the consensus sequences of TEs in *D. melanogaster* (Quesneville et al., 2005) and estimated the normalized abundance (reads per million) of the TEs in these two strains with our novel tool DeviaTE (Weilguny and Kofler, 2019). Apart from the telomeric TEs (TART-A, TART-B and TAHRE) which show distinct evolutionary dynamics (Pardue and DeBaryshe, 2011; Saint-Leandre and Levine, 2020), the most striking difference between the two strains was due to the LTR retrotransposon Tirant (fig. 1A). As expected, hobo and the I-element, two TEs that invaded *D. melanogaster* recently, are more abundant in the Iso-1 strain than in the older Canton-S strain (fig. The P-element is not present in both strains. To further investigate the abundance of Tirant in the two strains, we calculated the coverage of reads along the Tirant sequence with DeviaTE (fig. 1B; (Weilguny and Kofler, 2019)). We observed striking coverage differences between Canton-S and Iso-1 over the entire sequence of Tirant (fig. 1B; average normalized coverage; Iso-1=20.9, Canton-S=0.86;). Only few highly diverged reads aligned to Tirant in Canton-S (fig. 1B). In addition to these diverged reads, many reads with a high similarity to the consensus sequence of Tirant aligned in Iso-1 (fig. 1B).

**Figure 1:**
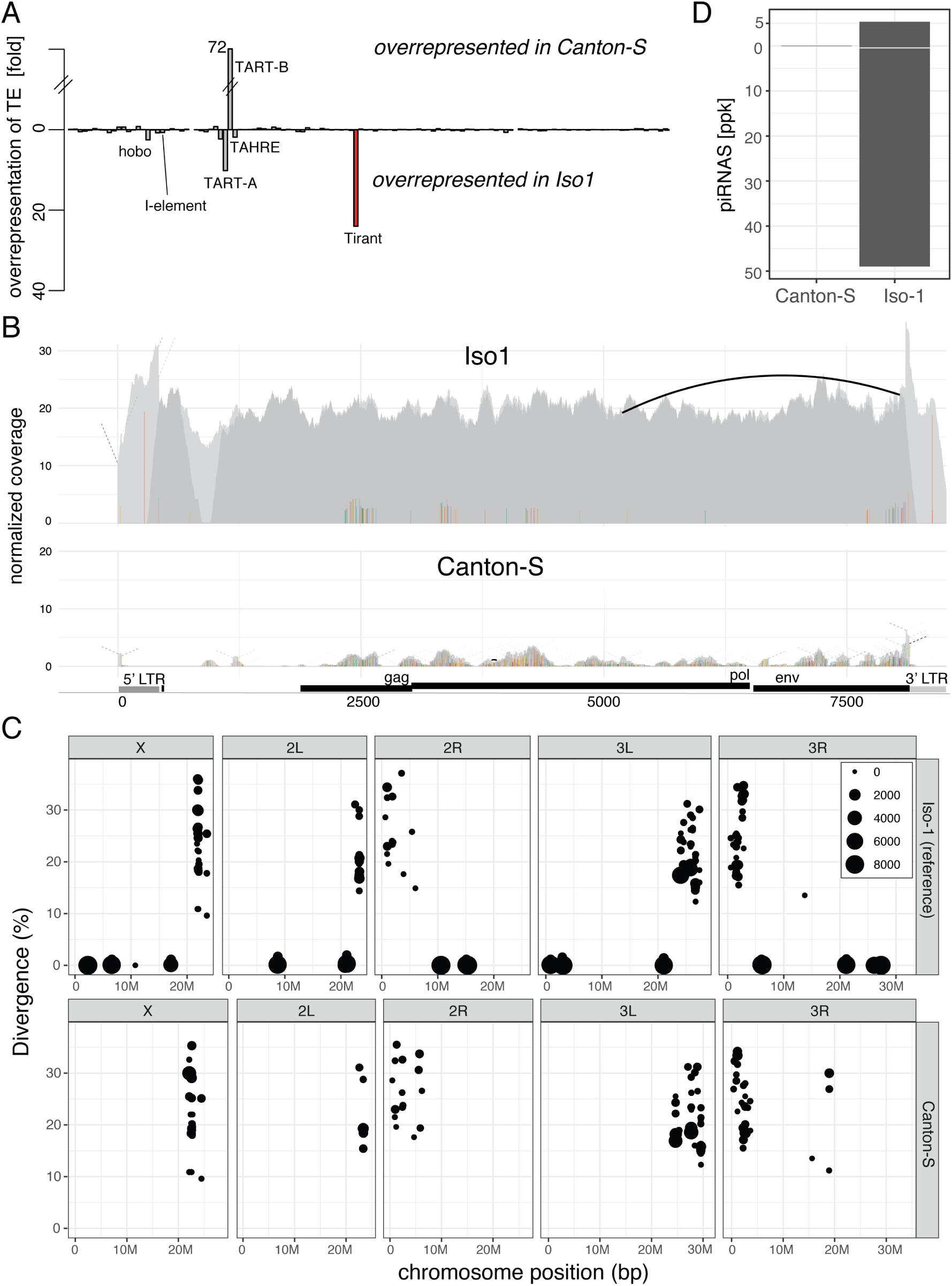
Canonical Tirant insertions are present in Iso-1 but not Canton-S. A) Differences in TE content between Canton-S and Iso-1. For each TE family (x-axis), we show the fold-difference in the number of reads mapping to a TE (y-axis) between the two strains. Note that reads mapping to Tirant (red) are overrepresented in Iso-1. B) Abundance and diversity of Tirant in Iso-1 and Canton-S. Short reads were aligned to the consensus sequence of Tirant and visualized with DeviaTE. The coverage of Tirant was normalized to the coverage of single copy genes. Single nucleotide polymorphisms (SNPs) and small internal deletions (indels) are shown as colored lines. Large internal deletions are shown as black arcs. Note that solely a few, highly degraded copies of Tirant are present in Canton-S C) Overview of Tirant insertions in the genomes of Iso-1 and Canton-S. For each Tirant insertion we show the position in the assembly, the length (size of dot), and the similarity to the consensus sequence (divergence). D) Abundance of sense (+ axis) and antisense (-axis) piRNAs for Tirant. ppk, piRNAs per 1000 miRNAs.

We refer to Tirant sequences with a high similarity to the consensus sequence as “canonical” Tirant. To identify the genomic location of the canonical and the diverged Tirant sequences, we annotated TEs in publicly available assemblies of Canton-S (based on Oxford Nanopore long-read data) and Iso-1 (i.e. the reference genome) with RepeatMasker (fig. 1C; (Hoskins et al., 2015; Wierzbicki et al., 2020)). Both assemblies are of high quality and suitable for genomic analysis of TEs (Wierzbicki et al., 2020). In Canton-S, only highly fragmented and diverged Tirant sequences were found close to the centromeres (fig. 1C). In addition to these diverged Tirant sequences, Iso-1 carries several canonical Tirant insertions on each chromosome arm (fig. 1C). This genomic distribution of Tirant, i.e. degraded Tirant fragments in the heterochromatin and canonical insertions in the euchromatin of *D. melanogaster*, was also noted in previous studies (Marsano et al., 2000; Mugnier et al., 2008). The absence of canonical Tirant insertions in euchromatin is also found in an independent assembly of Canton-S which is based on PacBio reads (supplementary fig. 1; (Chakraborty et al., 2019)). It was proposed that the degraded Tirant insertions located in heterochromatin may be due to an ancient invasion of Tirant, predating the split of *D. melanogaster* and *D. simulans* (Fablet et al., 2007; Lerat et al., 2011). It was further proposed that canonical insertions are of more recent origin (Bowen and McDonald, 2001; Lerat et al., 2011; Rahman et al., 2015). We thus speculated that the canonical insertions of Tirant may have recently been active while the degraded insertions in the heterochromatic may be inactive for some time (see also (Mugnier et al., 2008; Fablet et al., 2009)). If this is true, canonical insertions ought to segregate at low frequency in natural populations while the degraded insertions should mostly be fixed. To test this hypothesis, we estimated the population frequencies of the canonical and the degraded Tirant insertions in a natural *D. melanogaster* population from France (Viltain) (Kapun et al., 2018) with PoPoolationTE2 (Kofler et al., 2016). Indeed, most canonical Tirant insertions segregate at a low population frequency (*f* = 0.063) in the euchromatin, whereas most degraded insertions are in the heterochromatin and segregate at significantly higher frequencies (*f* = 0.73; Wilcoxon Rank sum test *p <* 2.2*e -* 16; supplementary fig. 2). Due to relaxed purifying selection in low-recombining regions (Eanes et al., 1992; Sniegowski and Charlesworth, 1994; Bartolomé et al., 2002; Petrov et al., 2011; Kofler et al., 2012), degraded Tirant insertions may have accumulated in the heterochromatin. Taken together, we hypothesize that Tirant invaded natural *D. melanogaster* populations in at least two waves of activity: an ancient wave, possibly predating the split of *D. melanogaster* and *D. simulans*, and a recent wave after Canton-S was sampled.

If Tirant recently invaded *D. melanogaster* populations, we expect to see differences in the composition of piRNAs between Iso-1 and Canton-S. Strains invaded by Tirant, such as Iso-1, should have established a functional defense against the TE and thus generate large amounts of piRNAs complementary to canonical Tirant. Contrarily, naive strains, such as Canton-S, should have few Tirant piRNAs. To test this, we sequenced piRNAs from the ovaries of both strains. Indeed, piRNAs against Tirant were highly abundant in Iso-1 but not in Canton-S (fig. 1D). Compared to the piRNA abundance of other TE families in *D. melanogaster*, Tirant piRNAs rank amongst the most abundant in Iso-1 but the least abundant in Canton-S (supplementary fig. 3A). Both sense and antisense piRNAs are distributed over the entire sequence of Tirant in Iso-1 (supplementary fig. 3B). TEs that are silenced in the germline by dual-strand clusters show a characteristic 10nt overlap between sense and antisense piRNAs, i.e. the ping-pong signature (Brennecke et al., 2007; Malone et al., 2009). Tirant has a pronounced ping-pong signature in Iso-1 but not in Canton-S (supplementary fig. 3C). We thus conclude that a piRNA-based defense mechanism against Tirant is active in Iso-1 but not in Canton-S.

If Tirant invaded natural *D. melanogaster* populations recently, old strains should only have a few highly degraded Tirant sequences (similar to Canton-S), whereas more recently collected strains should have many insertions with a high similarity to the consensus sequence of Tirant (i.e. canonical Tirant insertions). To test this, we sequenced 12 of the oldest available *D. melanogaster* strains (sampled between 1920 and 1970; fig. 2; supplementary table 1). Additionally, we included publicly available data of 15 different *D. melanogaster* strains into the analyses (fig. 2A; supplementary table 1). The reads were mapped to the consensus sequences of TEs in *Drosophila* and the TE abundance was assessed with DeviaTE (supplementary fig 4; (Weilguny and Kofler, 2019)). Strikingly, six out of seven strains sampled before or in 1938 solely contained degraded Tirant sequences (supplementary table 1; supplementary fig 4). The first strain carrying canonical Tirant sequences (Urbana-S) was collected around 1938. All 16 strains collected around or after 1950 carried canonical Tirant sequences (supplementary table 1). Estimates of the TE copy numbers support these observations (fig. 2A). Our results thus suggest that the canonical Tirant invaded *D. melanogaster* populations between 1938 and 1950 (fig. 2), whereas the degraded Tirant sequences are of more ancient origin, possibly predating the split of *D. melanogaster* and *D. simulans* (Fablet et al., 2007; Lerat et al., 2011). Since we were interested in the timing of the Tirant invasion relative to the other three TEs that recently invaded *D. melanogaster* populations, we also investigated the abundance and diversity of the I-element, hobo and the P-element in these strains (supplementary table 1; supplementary fig. 5, 6, 7). Interestingly, our data suggest that Tirant invaded natural *D. melanogaster* populations just before the I-element, followed by hobo and, lastly, by the P-element (supplementary table 1; fig. 2B).

**Figure 2:**
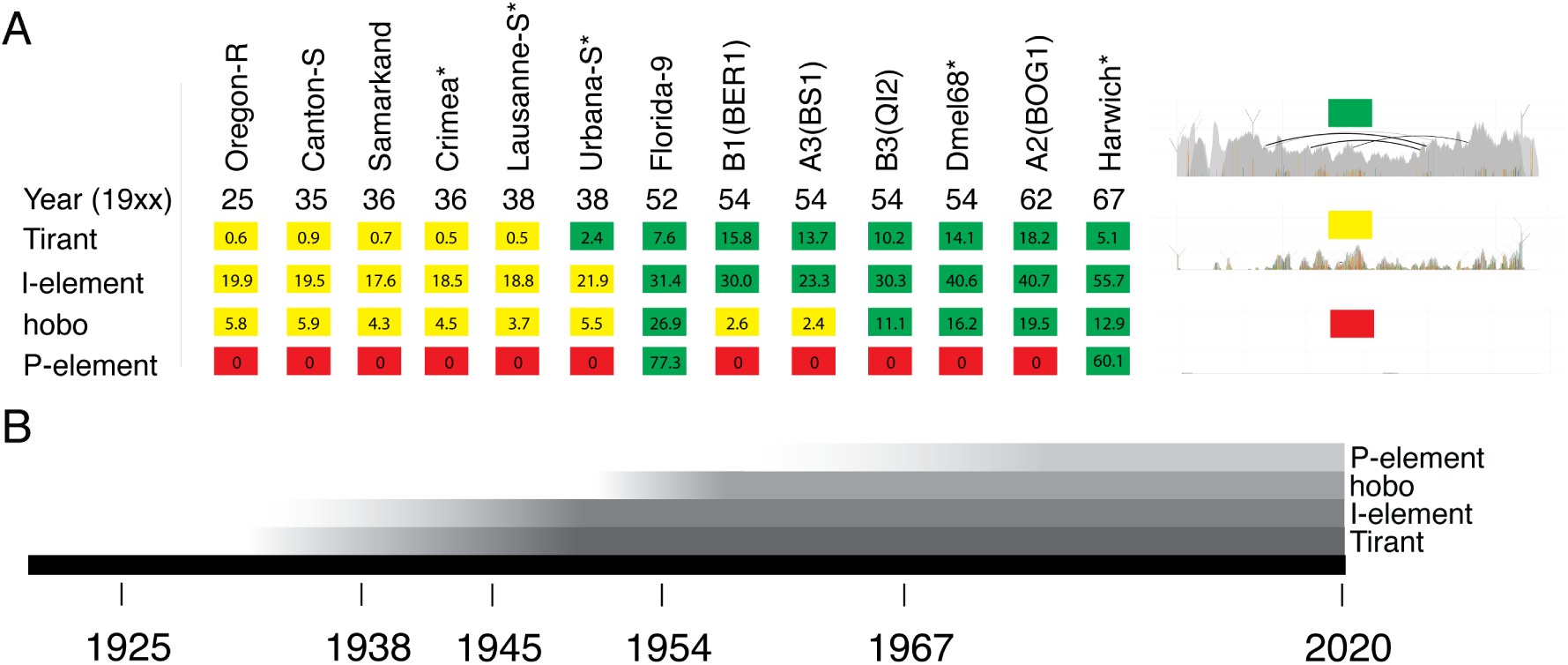
History of recent TE invasions in natural *D. melanogaster* populations. A) Overview of Tirant, I-element, hobo, and P-element sequences in some *D. melanogaster* strains. For an overview of these TEs in all investigated strains see supplementary table 1. The year corresponds to the sampling date of the strain. For each family, we classified the TE content into three distinct categories: red, absence of any TE sequence; yellow, solely degraded TE sequences are present; green, non-degraded sequences, with a high similarity to the consensus sequence are present; Numbers represent estimates of TE copy numbers per haploid genome obtained with DeviaTE. The abundance of degraded copies may be underestimated as copy-number estimates are based on the average coverage of the consensus sequence. The right panel shows an example for each of the three categories (similar to fig. 1B). B) Timeline showing the estimated invasion history of Tirant, the I-element, hobo and the P-element.

To further investigate the Tirant composition among strains, we performed a PCA based on the allele frequencies of Tirant single nucleotide polymorphism (SNPs) (fig. 3). Note that these allele frequency estimates solely reflect the Tirant composition within a particular strain (e.g. if 14 Tirant insertions in a given strain carry an ‘A’ at some site and 6 a ‘T’, the frequency of ‘A’ at this site is 0.7). In addition to the above mentioned strains (supplementary table 1), we also analyzed the Tirant content of natural populations. To do this, we relied on the Global Diversity Lines (GDL), i.e. several *D. melanogaster* strains sampled after 1988 (Begun and Aquadro, 1995) from five different continents (Africa - Zimbabwe, Asia - Beijing, Australia - Tasmania, Europe - Netherlands, America - Ithaca; (Grenier et al., 2015)).

**Figure 3:**
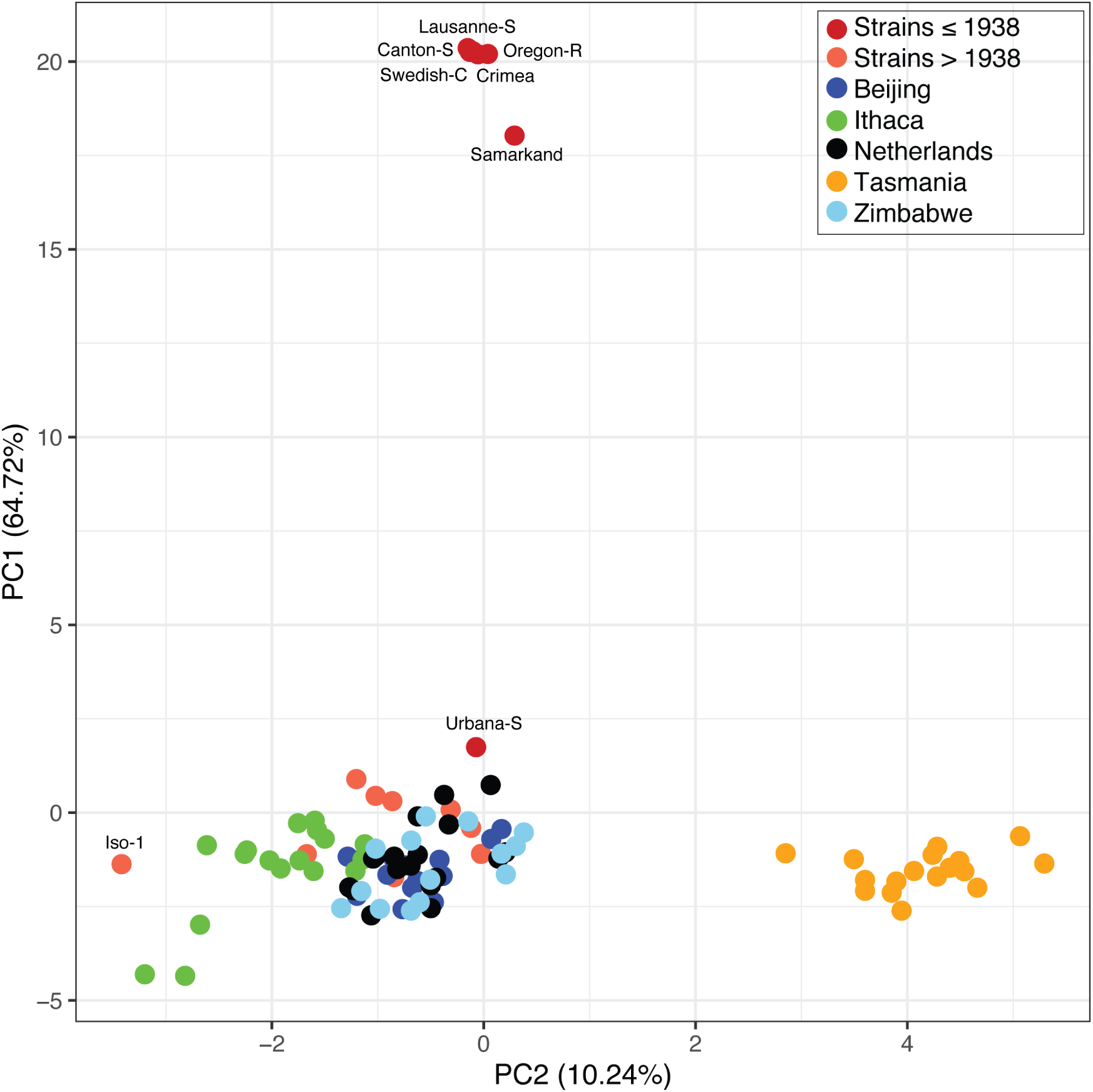
PCA based on the allele frequencies of Tirant SNPs in different *D. melanogaster* strains. In addition to previously described *D. melanogaster* strains, the Global Diversity Lines (GDL) were analysed. Note that the strains without canonical Tirant insertions as well as populations from Tasmania form distinct groups.

Old strains, collected before 1938, formed a distinct group (fig. 3), supporting our view that they carry distinct Tirant sequences. By contrast, most strains collected after 1938 and the majority of the GDLs group into one large cluster (fig. 3). All GDL strains thus carry non-degraded Tirant sequences. This observation also holds when additional, recently collected *D. melanogaster* strains are analyzed (e.g. DGRP, DrosEU, DrosRTEC; supplementary fig. 10; (Mackay et al., 2012; Bergland et al., 2014; Lack et al., 2015; Kapun et al., 2018; Machado et al., 2019)). Our data thus suggest that Tirant invaded most worldwide *D. melanogaster* populations. The reference strain Iso-1 is distant to the large cluster (fig. 3). Closer inspection revealed that Tirant insertions from natural populations carry 8 SNPs that are not found in the reference strain (supplementary fig. 8; supplementary table 2). Interestingly, also strains collected from Tasmania (Australia) formed a distinct group (fig. 3). This is due to multiple SNPs having markedly different allele frequencies in Tasmanian populations than in populations from other geographic locations (supplementary fig. 9; supplementary table 3). For hobo and the I-element, Tasmanian populations did not form a separate cluster (supplementary fig. 11; the P-element is absent in many samples, hence allele frequencies could not be calculated). This raises the question of what processes could be responsible for such striking differences in the Tirant composition among natural populations? We suggest that the Tirant invasion in Tasmania was subject to a founder effect, where flies carrying some rare variants of Tirant migrated to Tasmania, thereby triggering the spread of these rare Tirant variants in Tasmanian populations. Similarly, the strains used for generating Iso-1 may have carried rare Tirant variants that multiplied in these lines after they were sampled. In agreement with this, most Iso-1 specific SNPs segregate at low frequency in some *D. melanogaster* populations from Europe and North America (supplementary table 8).

In summary, we conclude that Tirant invaded all investigated worldwide populations of *D. melanogaster* during the past century. Furthermore, founder effects may be important components of TE invasions, since they may lead to a geographically heterogenous TE composition.

The other three TEs that invaded *D. melanogaster* populations within the last 100 years caused some hybrid dysgenesis (HD) symptoms. To test whether Tirant also induces HD symptoms, we performed crosses between a strain having recent Tirant insertions (Urbana-S) and a strain not having such insertions (Lausanne-S). Both strains do not have recent P-element, I-element and hobo insertions, which rules out interference by the other HD systems (fig. 2A; supplementary table 1). We investigated the fraction of dysgenic ovaries in the F1 generation, a trait influenced by P-element and hobo mobilization (Kidwell et al., 1977; Yannopoulos et al., 1987; Blackman et al., 1987), and the fraction of hatched F2 embryos, a trait influenced by I-element mobilization (Bucheton et al., 1976). We performed all crosses at several temperatures (see methods), as temperature frequently has a strong influence on the extent of HD symptoms (Kidwell et al., 1977; Bucheton, 1979; Kidwell and Novy, 1979; Serrato-Capuchina et al., 2020). We did not find any significant differences in the number of dysgenic ovaries nor in the number of hatched eggs between the reciprocal crosses (supplementary fig. 12; supplementary table 4). We hypothesize that the absence of apparent HD symptoms may be one reason why the spread of Tirant in natural *D. melanogaster* populations during the past century was not detected before.

Lastly, we aimed to shed light on the origin of the Tirant invasion. Since canonical Tirant insertions are mostly absent in strains collected before 1938, we reasoned that the recent Tirant invasion was likely triggered by HT (or an introgression). To identify the putative donor species, we investigated Tirant sequences in different *Drosophila* species. We first tested if Tirant sequences can be found in 11 sequenced *Drosophila* genomes (Drosophila 12 Genomes Consortium, 2007). Solely members of the *Drosophila melanogaster* species subgroup contained reads mapping to Tirant (supplementary fig. 13; *D. melanogaster, D. simulans, D. erecta, D. yakuba*). To further investigate the composition of Tirant in the *melanogaster* subgroup, we obtained Illumina short read data for several individuals from different species of this subgroup. In addition to *D. melanogaster, D. simulans, D. erecta* and *D. yakuba* we also obtained data for *D. sechellia, D. mauritiana* and *D. teisseri* (supplementary table 5). A PCA based on Tirant SNPs showed that recently collected *D. melanogaster* strains (*>*1938) cluster close to *D. simulans* strains (supplementary fig. 14). The high similarity of Tirant sequence between *D. melanogaster* and *D. simulans* was noted before (Fablet et al., 2006, 2007; Lerat et al., 2011; Bargues and Lerat, 2017). This raises the possibility that a HT from *D. simulans* triggered the recent Tirant invasion in *D. melanogaster*. Interestingly, Tirant sequences of old laboratory strains collected before 1938, cluster with Tirant sequences of *D. erecta*, which also suggests that the degraded Tirant sequences are ancient, possibly predating the split of the melanogaster subgroup (Fablet et al., 2007; Lerat et al., 2011).

## Discussion

We showed that the retrotranspon Tirant invaded natural *D. melanogaster* populations during the last 100 years. This conclusion is based on the absence of canonical Tirant sequences in most strains collected before 1938 and their presence in strains collected after 1938. As an alternative explanation, most strains collected before 1938 could have lost the canonical Tirant sequences. It was, for example, proposed that non-African *D. simulans* populations lost canonical Tirant sequences (Fablet et al., 2006). But this alternative explanation seems unlikely as it requires the independent loss of canonical Tirant sequences in strains collected before 1938 but not in any strain collected after 1938. The low population frequency of euchromatic Tirant insertions (see also Kofler et al., 2015b) and the high sequence similarity between the left and the right LTR of Tirant insertions (Bowen and McDonald, 2001) are also in agreement with our hypothesis of a recent Tirant invasion (both LTRs are identical upon insertion of a TE but mutate over time). Our hypothesis of the recent Tirant invasion is also consistent with the interpretation of the data for the I-element, P-element and hobo, where the absence of the (canonical) TE in old strains combined with the presence in young strains was taken as evidence for recent invasions (Kidwell, 1983; Daniels et al., 1990a,b; Bucheton et al., 1992).

Our data suggest that Tirant was the first TE that invaded natural *D. melanogaster* populations in the last century. However, these results need to be interpreted with caution as i) there is some uncertainty about the sampling time of the strains, ii) some strains may have been contaminated (e.g. the presence of the P-element in a strain collected around 1938 (Swedish-C) is likely due to contamination; supplementary table 1), and iii) our strains are from different geographic regions, where some regions might have been invaded earlier than others. Nevertheless, our results are largely in agreement with previous works which suggested that the I-element invasion happened between 1930 and 1950, the hobo invasion around 1955 and the P-element invasion between 1950 and 1980 (Kidwell, 1983; Periquet et al., 1989; Anxolabéhère et al., 1988).

We did not find evidence that Tirant induces HD symptoms. Also, a previous work in *D. simulans* did not report HD symptoms for Tirant despite Tirant being activated by reciprocal crosses (Akkouche et al., 2013). However, due to several reasons, more work will be necessary to show whether Tirant causes some HD symptoms. First, it is not clear what symptoms to look for. We investigated the fraction of dysgenic ovaries in the F1 and the fraction of hatched eggs (F2), two traits classically affected by HD. However, it is feasible that Tirant activity leads to entirely different phenotypic effects, especially since Tirant might be an endogenous retrovirus that multiplies by a different mechanism than the other three TEs that cause HD symptoms; Tirant may infect the germline via virus-like-particles, whereas the other three TEs are directly active in the germline (Calvi and Gelbart, 1994; Akkouche et al., 2012; Wang et al., 2018; Moon et al., 2018; Barckmann et al., 2018). Second, the severity of HD symptoms frequently depends on multiple factors, such as temperature and the age of flies (Kidwell et al., 1977; Bucheton, 1979; Kidwell and Novy, 1979; Serrato-Capuchina et al., 2020). It is feasible that HD symptoms of Tirant can only be observed under certain conditions. Third, previous studies noted marked differences in the ability to induce or repress HD among different strains (Kidwell et al., 1977, 1988; Anxolabéhère et al., 1988; Srivastav et al., 2019). It is thus feasible that solely crosses of certain strains show HD symptoms for Tirant.

It is currently unclear how canonical Tirant sequences entered *D. melanogaster* populations. Possible explanations are HT or introgression from a related species (Silva et al., 2004; Sánchez-Gracia et al., 2005; Loreto et al., 2008; Bartolomé et al., 2009). In search for a possible donor species we found that the Tirant sequences of *D. melanogaster* and *D. simulans* are closely related (see also (Fablet et al., 2007; Lerat et al., 2011; Bargues and Lerat, 2017)). Non-degraded Tirant sequences in *D. simulans* (Tirant-C; Fablet et al. (2006)) are closer related to the canonical Tirant sequences in *D. melanogaster* than to the degraded Tirant sequences of *D. simulans* (Tirant-S) (Lerat et al., 2011; Bargues and Lerat, 2017). This indicates a close phylogenetic relationship of Tirant (C type) in *D. simulans* and *D. melanogaster*, possibly exceeding the similarity expected with vertical transfer of Tirant (Lerat et al., 2011). HT of TEs between these two species is plausible since both species are closely related (Lemeunier et al., 1976) and have largely overlapping habitats (Parsons and Stanley, 1981), which generates ample opportunities for HT or introgressions. HT of TEs between these species was observed before in both directions. For example, Kofler et al. (2015b) suggested that *D. simulans* recently acquired the P-element from *D. melanogaster*. Conversely, hobo and the I-element in *D. melanogaster* were possibly acquired from *D. simulans* (Daniels et al., 1990a; Simmons, 1992; Loreto et al., 2008).

We found that Tirant sequences from Tasmania (an island south of Australia) have a different composition than Tirant sequences from other locations (at least 5 SNPs have distinctly different frequencies; supplementary table 3). We suggest that this may be due to a founder effect during the Tirant invasion, which led to the spread of rare Tirant variants in Tasmanian populations. We wondered whether the observed founder effect could be due to the recent colonization of Australia (Tasmania) by D. melanogaster (Bock and Parsons, 1981). However, this seems unlikely as the colonization of Australia, and probably also of Tasmania, predates the Tirant invasion. *D. melanogaster* was first spotted in Australia in 1894 and is known to rapidly spread into nearby areas (Bock and Parsons, 1981; Keller, 2007), whereas the Tirant invasion likely happened between 1938 and 1950. Moreover, founder effects that occurred during the colonization of Tasmania should affect the entire genomic background of *D. melanogaster* and not just the Tirant sequences. Previous studies did not detect any signatures of bottlenecks for Tasmanian *D. melanogaster* populations (Agis and Schlötterer, 2001; Grenier et al., 2015; Bergland et al., 2016; Arguello et al., 2019). We thus argue that the founder effect in Tasmania is specific to Tirant. Founder effects during TE invasions could be important, hitherto little considered, processes that may lead to geographically distinct TE compositions.

We suggest that four different TEs invaded *D. melanogaster* populations within 40 years (between the 1930s and 1970s). Why did so many different TEs spread in *D. melanogaster* within such a short time? A possible explanation could be the recent habitat expansion of *D. melanogaster* into the Americas and Australia about 100-200 years ago (Bock and Parsons, 1981; Vieira et al., 1999; Kofler et al., 2015b). Habitat expansion may bring species into contact that did not co-exist before. If these species carry different TE families, HT events between the species may trigger novel TE invasions. A classic example is the P-element in *D. melanogaster* which was likely acquired from *D. willistoni* after *D. melanogaster* entered the habitat of *D. willistoni* in South America (Engels, 1992). The lag-time between colonization of the Americas and Australia (about 100-200 years ago; (Bock and Parsons, 1981; Keller, 2007)) and the four different TE invasions (1930-1970) may be due to the stochasticity of HT events, a strong influence of drift in the early stages of TE invasions and the time required until a TE reaches an appreciable frequency (Ginzburg et al., 1984; Le Rouzic and Capy, 2005). It will be interesting to see if such a high rate of novel TE invasions in *D. melanogaster* populations will be maintained over the next century. An absence of novel invasions would support our hypothesis that the habitat expansion triggered the four recent TE invasions in *D. melanogaster*.

Out of the four TEs that invaded *D. melanogaster* populations in the last century, the P-element is unique as it is the only TE that does not show similarity to any sequence of the *D. melanogaster* genome. For the other three TEs - Tirant, the I-element and hobo - many degraded insertions can be found (mostly in the heterochromatin) (Bucheton et al., 1984, 1986; Daniels et al., 1990a; Bucheton et al., 1992)), likely due to previous invasions. Thus, three out of the four TEs probably invaded *D. melanogaster* populations at least twice. This raises the question of how multiple waves of invasions arise. Before a TE can trigger a novel invasion the TE needs to overcome the host defense (or the host defense may break down). For example, in mammals and invertebrates efficient silencing of a TE requires piRNAs that match with less than 10% sequence divergence over the bulk of the TE sequence (Post et al., 2014; Kotov et al., 2019). A TE that diverged by more than 10% from the piRNA pool of the host could thus trigger a second wave of an invasion. The same consideration holds for other host defence mechanism that rely on sequence similarity to a TE, like small RNAs in plants or KRAB-ZNFs in mammals (Yang et al., 2017; Marí-Ordóñez et al., 2013). It is however an important open question whether sufficient sequence divergence could be acquired within a host species, where host defence mechanisms may co-adapt with the TE, or whether HT to an intermediate host (e.g. a closely related species) is necessary to overcome the host defense.

## Materials and Methods

### Strains and dating

The sequenced fly strains were obtained from the Bloomington Drosophila Stock Center (BDSC) (Crimea, Lausanne-S, Swedish-C, Urbana-S, Berlin-K, Hikone-R, Florida-9, Pi2, Dmel68, Harwich, w1118, wk, Amherst-3). We additionally analysed publicly available sequencing data of different *D. melanogaster* strains (Grenier et al., 2015; King et al., 2012; Mackay et al., 2012; Bergland et al., 2014; Lack et al., 2015; Jakšić et al., 2017; Kapun et al., 2018; Machado et al., 2019; Wierzbicki et al., 2020) (supplementary table 5). The collection dates of the strains were obtained from different sources. If available we used the collection dates from Lindsley and Grell (1968). Alternatively, we used the collection dates published in previous works (Engels, 1979; Black et al., 1987; Anxolabéhère et al., 1988; Galindo et al., 1995; Ruebenbauer et al., 2008) or information from the National Drosophila Species Stock Center (drosophilaspecies.com) and FlyBase (flybase.org/reports/FBrf0222222.html) (supplementary table 1). For the strains w1118 and Urbana-S we used the latest possible collection date: for w1118 we used the publication date of the first publication mentioning the strain and for Urbana-S we used the year of the death of C. Bridges, who collected the strain (Lindsley and Grell, 1968) (supplementary table 1). The geographic origin was obtained from the same sources. For an overview of the used strains, the estimated collection date and the source of the information see supplementary table 1. Additionally we used publicly available data of different strains from *D. simulans, D. sechellia, D. mauritiana, D. yakuba, D. teisseri* and *D. erecta* (Garrigan et al., 2012, 2014; Rogers et al., 2014; Turissini et al., 2015; Miller et al., 2018; Schrider et al., 2018; Melvin et al., 2018; Lanno et al., 2019; Kang et al., 2019; Meany et al., 2019; Cooper et al., 2019; Stewart and Rogers, 2019; Drosophila 12 Genomes Consortium, 2007). For an overview of all used publicly available data see supplementary table 5.

### DNA Sequencing

DNA for Illumina paired-end sequencing was extracted from whole bodies of 20-30 virgin female flies using a salt-extraction protocol (Maniatis et al., 1982). Libraries were prepared with the NEBNext Ultra II DNA library Prep Kit (New England Bioloabs, Ipswich, MA) using 1 *µ*g DNA. Illumina sequencing was performed by the Vienna Biocenter Core Facilities using the HiSeq2500 platform (2×125bp; Illumina, San Diego, CA, USA).

### small RNA sequencing

For small RNA sequencing, we extracted total RNA from ovaries of the strains Canton-S and Iso-1 using TRIzol. The small RNA was sequenced by Fasteris (Geneva, Switzerland). After depletion of 2S rRNA, library preparation was performed using the Illumina TruSeq small RNA kit and cDNA was sequenced on an Illumina NextSeq platform (50bp; Illumina, San Diego, CA, USA). Adapter sequences were trimmed with cutadapt (v2.3) (Martin, 2011) (adapter: TGGAATTCTCGGGTGCCAAGGAACTCCAGTCACCATTTTATCTCGTATGC) and filtered for reads with a length between between 18 and 36nt. The reads were mapped to a database consisting of *D. melanogaster* miRNAs, mRNAs, rRNAs, snRNAs, snoRNAs, tRNAs (Thurmond et al., 2019) and the TE sequences (Quesneville et al., 2005) using novoalign (v3.09; http://novocraft.com/). Solely piRNAs with a length between 23 and 29nt were retained and the abundance of piRNAs was normalized to a million miRNAs as described previously (Kofler et al., 2018). For computing the ping-pong signatures and visualizing the piRNA abundance along the Tirant sequence we used a previously developed pipeline (Kofler et al., 2018).

### TE abundance and diversity

The coverage along a TE and the frequencies of SNPs and indels in a TE were computed using our newly developed tool DeviaTE (Weilguny and Kofler, 2019). Briefly short reads from a sample were aligned with bwa sw (v0.7.17) (Li and Durbin, 2010) to the TE consensus sequences of *Drosophila* (Quesneville et al., 2005) as well as to three single copy genes *(traffic jam, rpl32* and *rhino*), which allowed us to infer TE copy numbers by contrasting the coverage of a TE to the coverage of the single copy genes. The abundance and diversity of TE insertions was visualized with DeviaTE (Weilguny and Kofler, 2019). Based on these visualizations we manually classified the presence/absence of Tirant, hobo, the I-element and the P-element in different *D. melanogaster* strains (supplementary table 1). We used the following three categories: i) absence of any TE sequences, ii) solely degraded TE sequences are present, iii) non-degraded sequences, with a high similarity to the consensus sequence, are present. For examples see supplementary figures 4,5,6,7. Note that a PCA based on the allele frequencies of SNPs in a TE supports our classification for Tirant and hobo. Since many strains do not contain any P-element sequences, the allele frequencies of SNPs in the P-element could not be calculated for all strains. Despite discernible differences between strains with and without recent I-element insertions, the PCA did not separate these two groups (supplementary fig. 5,11). The PCA was performed in R (prcomp) using arcsine and square root transformed allele frequencies of SNPs in TEs (R Core Team, 2012). The DSPR lines were not included into the PCA due to their short read length (50bp).

Tirant sequences in the assemblies of Canton-S (Wierzbicki et al., 2020) and Iso-1 (v6.22; https://flybase.org/) were identified with RepeatMasker (open-4.0.7; Smit et al. (2015)) using the TE consensus sequences of *Drosophila* as custom library (Quesneville et al., 2005).

To estimate the position and population frequency of canonical and degraded Tirant insertions in a natural *D. melanogaster* population we used PoPoolationTE2 (Kofler et al., 2016) and a population collected in 2014 at Viltain (France) by the DrosEU consortium (SRR5647729; (Kapun et al., 2018)). We first extracted the sequences of degraded

Tirant insertions (*>* 10% divergence to consensus sequence) from the reference assembly (v6.22) with RepeatMasker and bedtools (Quinlan and Hall, 2010) (v2.29.2).Only degraded copies longer than 100bp were retained. We generated the artificial reference genome required by PoPoolationTE2, by merging the repeat masked reference genome, the consensus sequence of Tirant and the degraded Tirant sequences into a single fasta file. The short reads were mapped to this artificial reference genome using bwa mem (Li and Durbin, 2009) with paired-end mode and the parameter -M. The mapped reads were sorted with samtools (Li et al., 2009). Finally we followed the PoPoolationTE2 pipeline using the parameters: –map-qual 15, –min-count 2, –min-coverage 2. We indicated heterochromatic regions following previous work (Riddle et al., 2011; Hoskins et al., 2015).

### Hybrid dysgenesis assay

To test whether Tirant induces HD symptoms we performed reciprocal crosses between the *D. melanogaster* strains Urbana-S (having recent Tirant insertions) and Lausanne-S (not-having recent Tirant insertions). The choice of the two strains was based on the absence of recent I-element, hobo and P-element insertions in both strains, which avoids interference with other HD systems (supplementary table 1). Each cross was performed in 3 replicates by mating 20 female virgin flies with 15 males. To estimate the number of dysgenic ovaries, 2-5 days old F1 flies (kept at either 20 °C, 25 °Cor 29 °C) were allowed to lay eggs on black agar plates (containing charcoal) for 24 hours. The F1 female ovaries were dissected on PBS and scored for the presence of dysgenic (underdeveloped) ovaries. The deposited F2 embryos were counted, incubated for 24 hours and the number of larvae (=hatched eggs) was quantified.

## Author contributions

FS and RK conceived the work. FS and FW analyzed the data. KS provided feedback on the manuscript. FS and RK wrote the manuscript.

## Acknowledgements

We thank Elisabeth Salbaba for technical support and Divya Selvaraju for providing the Harwich data. We thank all members of the Institute of Population Genetics for feedback and support. This work was supported by the Austrian Science Foundation (FWF) grants P30036-B25 to RK and W1225.

## Data availability

All scripts are available at https://sourceforge.net/projects/te-tools/ (folder tirant) and important files (including all DeviaTE outputs) at https://sourceforge.net/projects/te-tools/files/tirant_data/. The sequence data of the old laboratory strains and the piRNA sequences are available at NCBI (PRJNA634847).

## References

Agis, M. and Schlötterer, C. (2001). Microsatellite variation in natural *Drosophila melanogaster* populations from New South Wales (Australia) and Tasmania. Molecular Ecology, 10(5):1197–1205.

Akkouche, A., Grentzinger, T., Fablet, M., Armenise, C., Burlet, N., Braman, V., Chambeyron, S., and Vieira, C. (2013). Maternally deposited germline piRNAs silence the tirant retrotransposon in somatic cells. EMBO Reports, 14(5):458–464.

Akkouche, A., Rebollo, R., Burlet, N., Esnault, C., Martinez, S., Viginier, B., Terzian, C., Vieira, C., and Fablet, M. (2012). tirant, a Newly Discovered Active Endogenous Retrovirus in *Drosophila simulans*. Journal of Virology, 86(7):3675–3681.

Anxolabéhère, D., Kidwell, M. G., and Periquet, G. (1988). Molecular characteristics of diverse populations are consistent with the hypothesis of a recent invasion of *Drosophila melanogaster* by mobile P elements. Molecular Biology and Evolution, 5(3):252–269.

Arguello, J. R., Laurent, S., Clark, A. G., and Gaut, B. (2019). Demographic History of the Human Commensal *Drosophila melanogaster*. Genome Biology and Evolution, 11(3):844–854.

Barckmann, B., El-Barouk, M., Pélisson, A., Mugat, B., Li, B., Franckhauser, C., Fiston Lavier, A.-S., Mirouze, M., Fablet, M., and Chambeyron, S. (2018). The somatic piRNA pathway controls germline transposition over generations. Nucleic Acids Research, 46(18):1–13.

Bargues, N. and Lerat, E. (2017). Evolutionary history of LTR-retrotransposons among 20 *Drosophila* species. Mobile DNA, 8(1):1–15.

Bartolomé, C., Bello, X., and Maside, X. (2009). Widespread evidence for horizontal transfer of transposable elements across *Drosophila* genomes. Genome Biology, 10(2).

Bartolomé, C., Maside, X., and Charlesworth, B. (2002). On the abundance and distribution of transposable elements in the genome of *Drosophila melanogaster*. Molecular biology and evolution, 19(6):926–937.

Begun, D. J. and Aquadro, C. F. (1995). Molecular variation at the vermilion locus in geographically diverse populations of *Drosophila melanogaster* and *D. simulans*. Genetics, 140(3):1019–1032.

Bergland, A. O., Behrman, E. L., O’Brien, K. R., Schmidt, P. S., and Petrov, D. A. (2014). Genomic Evidence of Rapid and Stable Adaptive Oscillations over Seasonal Time Scales in *Drosophila*. PLoS Genetics, 10(11).

Bergland, A. O., Tobler, R., González, J., Schmidt, P., and Petrov, D. (2016). Secondary contact and local adaptation contribute to genome-wide patterns of clinal variation in *Drosophila melanogaster*. Molecular Ecology, 25(5):1157–1174.

Biémont, C. (2010). A brief history of the status of transposable elements: From junk DNA to major players in evolution. Genetics, 186(4):1085–1093.

Bingham, P. M., Kidwell, M. G., and Rubin, G. M. (1982). The molecular basis of P-M hybrid dysgenesis: The role of the P element, a P-strain-specific transposon family. Cell, 29(3):995–1004.

Black, D. M., Jackson, M. S., Kidwell, M. G., and Dover, G. A. (1987). KP elements repress P-induced hybrid dysgenesis in *Drosophila melanogaster*. The EMBO Journal, 613(13):4125–4135.

Blackman, R. K., Grimaila, R., Macy, M., Koehler, D., and Gelbart, W. M. (1987). Mobilization of hobo elements residing within the decapentaplegic gene complex: Suggestion of a new hybrid dysgenesis system in *Drosophila melanogaster*. Cell, 49(4):497–505.

Blumenstiel, J. P. (2011). Evolutionary dynamics of transposable elements in a small RNA world. Trends in Genetics, 27(1):23–31.

Bock, I. and Parsons, P. (1981). Species of Australia and New Zealand. In Ashburner, M., Carson, L., and Thompson, J. J., editors, The genetics and biology of Drosophila, volume 3a, pages 349–393. Academic Press, Oxford.

Bonnivard, E., Bazin, C., Denis, B., and Higuet, D. (2000). A scenario for the hobo transposable element invasion, deduced from the structure of natural populations of *Drosophila melanogaster* using tandem TPE repeats. Genetical Research, 75(1):13–23.

Bowen, N. J. and McDonald, J. F. (2001). *Drosophila* euchromatic LTR retrotransposons are much younger than the host species in which they reside. Genome Research, 11(9):1527–1540.

Brennecke, J., Aravin, A. A., Stark, A., Dus, M., Kellis, M., Sachidanandam, R., and Hannon, G. J. (2007). Discrete small RNA-generating loci as master regulators of transposon activity in *Drosophila*. Cell, 128(6):1089–1103.

Brennecke, J., Malone, C. D., Aravin, A. A., Sachidanandam, R., Stark, A., and Hannon, G. J. (2008). An epigenetic role for maternally inherited piRNAs in transposon silencing. Science, 322(5906):1387–1392.

Brizuela, B. J., Elfring, L., Ballard, J., Tamkun, J. W., and Kennison, J. A. (1994). Genetic analysis of the *brahma* gene of *Drosophila melanogaster* and polytene chromosome subdivisions 72AB. Genetics, 137(3):803–813.

Bucheton, A. (1979). Non-Mendelian female sterility in *Drosophila melanogaster*: Influence of aging and thermic treatments. III. Cumulative effects induced by these factors. Genetics, 93(1):131–142.

Bucheton, A., Lavige, J., Picard, G., and L’heritier, P. (1976). Non-mendelian female sterility in *Drosophila melanogaster*: quantitative variations in the efficiency of inducer and reactive strains. Heredity, 36(3):305–314.

Bucheton, A., Paro, R., Sang, H. M., Pelisson, A., and Finnegan, D. J. (1984). The molecular basis of I-R hybrid Dysgenesis in *Drosophila melanogaster*: Identification, cloning, and properties of the I factor. Cell, 38(1):153–163.

Bucheton, A., Simonelig, M., Vaury, C., and Crozatier, M. (1986). Sequences similar to the I transposable element involved in I-R hybrid dysgenesis in *D. melanogaster* occur in other *Drosophila* species. Nature, 322(6080):650–652.

Bucheton, A., Vaury, C., Chaboissier, M. C., Abad, P., Pélisson, A., and Simonelig, M. (1992). I elements and the *Drosophila* genome. Genetica, 86(1-3):175–90.

Calvi, B. R. and Gelbart, W. M. (1994). The basis for germline specificity of the hobo transposable element in *Drosophila melanogaster*. The EMBO Journal, 13(7):1636–44.

Cañizares, J., Grau, M., Paricio, N., and Moltó, M. D. (2000). Tirant is a new member of the gypsy family of retrotransposons in *Drosophila melanogaster*. Genome, 43(1):9–14.

Chakraborty, M., Emerson, J. J., Macdonald, S. J., and Long, A. D. (2019). Structural variants exhibit widespread allelic heterogeneity and shape variation in complex traits. Nature Communications, 10(1):419275.

Cooper, J. C., Guo, P., Bladen, J., and Phadnis, N. (2019). A triple-hybrid cross reveals a new hybrid incompatibility locus between *D. melanogaster* and *D. sechellia*. bioRxiv, page 590588.

Daniels, S. B., Chovnick, A., and Boussyy, I. A. (1990a). Distribution of hobo transposable elements in the genus *Drosophila*. Molecular Biology and Evolution, 7(6):589–606.

Daniels, S. B., Peterson, K. R., Strausbaugh, L. D., Kidwell, M. G., and Chovnick, A. (1990b). Evidence for horizontal transmission of the P transposable element between *Drosophila* species. Genetics, 124(2):339–355.

Doolittle, W. F. and Sapienza, C. (1980). Selfish genes, the phenotype paradigm and genome evolution. Nature, 284(5757):601–603.

Drosophila 12 Genomes Consortium (2007). Evolution of genes and genomes on the *Drosophila* phylogeny. Nature, 450(7167):203–18.

Eanes, W. F., Wesley, C., and Charlesworth, B. (1992). Accumulation of P elements in minority inversions in natural populations of *Drosophila melanogaster*. Genetical research, 59(1):1–9.

Engels, W. R. (1979). Hybrid dysgenesis in *Drosophila melanogaster*: Rules of inheritance of female sterility. Genetics Research, 89(5-6):407–424.

Engels, W. R. (1992). The origin of P elements in *Drosophila melanogaster*. BioEssays, 14(10):681–686.

Fablet, M., Lerat, E., Rebollo, R., Horard, B., Burlet, N., Martinez, S., Brasset, È., Gilson, E., Vaury, C., and Vieira, C. (2009). Genomic environment influences the dynamics of the tirant LTR retrotransposon in *Drosophila*. The FASEB Journal, 23(5):1482–1489.

Fablet, M., McDonald, J. F., Biémont, C., and Vieira, C. (2006). Ongoing loss of the tirant transposable element in natural populations of *Drosophila simulans*. Gene, 375(1-2):54–62.

Fablet, M., Souames, S., Biémont, C., and Vieira, C. (2007). Evolutionary pathways of the tirant LTR retrotransposon in the *Drosophila melanogaster* subgroup of species. Journal of Molecular Evolution, 64(4):438–447.

Galindo, M. I., Ladevèze, V., Lemeunier, F., Kalmes, R., Periquet, G., and Pascual, L. (1995). Spread of the autonomous transposable element hobo in the genome of *Drosophila melanogaster*. Molecular Biology and Evolution, 12(5):723–734.

Garrigan, D., Kingan, S. B., Geneva, A. J., Andolfatto, P., Clark, A. G., Thornton, K. R., and Presgraves, D. C. (2012). Genome sequencing reveals complex speciation in the *Drosophila simulans* clade. Genome Research, 22(8):1499–1511.

Garrigan, D., Kingan, S. B., Geneva, A. J., Vedanayagam, J. P., and Presgraves, D. C. (2014). Genome diversity and divergence in *Drosophila mauritiana*: Multiple signatures of faster X evolution. Genome Biology and Evolution, 6(9):2444–2458.

Ginzburg, L. R., Bingham, P. M., and Yoo, S. (1984). On the theory of speciation induced by transposable elements. Genetics, 107(2):331–341.

Goodier, J. L. (2016). Restricting retrotransposons: A review. Mobile DNA, 7(16).

Grenier, J. K., Roman Arguello, J., Moreira, M. C., Gottipati, S., Mohammed, J., Hackett, S. R., Boughton, R., Greenberg, A. J., and Clark, A. G. (2015). Global diversity lines-a five-continent reference panel of sequenced *Drosophila melanogaster* strains. G3: Genes, Genomes, Genetics, 5(4):593–603.

Gunawardane, L. S., Saito, K., Nishida, K. M., Miyoshi, K., Kawamura, Y., Nagami, T., Siomi, H., and Siomi, M. C. (2007). A slicer-mediated mechanism for repeat-associated siRNA 5’ end formation in *Drosophila*. Science, 315(5818):1587–1590.

Hickey, D. A. (1982). Selfish DNA: A sexually transmitted nuclear parasite. Genetics, 101(3-4):519–531.

Hoskins, R. A., Carlson, J. W., Wan, K. H., Park, S., Mendez, I., Galle, S. E., Booth, B. W., Pfeiffer, B. D., George, R. A., Svirskas, R., et al. (2015). The Release 6 reference sequence of the *Drosophila melanogaster* genome. Genome research, 25(3):445–458.

Jakšić, A. M., Kofler, R., and Schlötterer, C. (2017). Regulation of transposable elements: Interplay between TE-encoded regulatory sequences and host-specific trans-acting factors in *Drosophila melanogaster*. Molecular Ecology, 26(19):5149–5159.

Kang, L., Rashkovetsky, E., Michalak, K., Garner, H. R., Mahaney, J. E., Rzigalinski, B. A., Korol, A., Nevo, E., and Michalak, P. (2019). Genomic divergence and adaptive convergence in *Drosophila simulans* from Evolution Canyon, Israel. Proceedings of the National Academy of Sciences, 116(24):11839 – 11844.

Kapun, M., Aduriz, M. G. B., Staubach, F., Vieira, J., Obbard, D., Goubert, C., Stabelli, O. R., Kankare, M., Haudry, A., Wiberg, R. A. W., et al. (2018). Genomic analysis of European *Drosophila melanogaster* populations on a dense spatial scale reveals longitudinal population structure and continent-wide selection. bioRxiv, page 313759.

Keller, A. (2007). *Drosophila melanogaster*’s history as a human commensal. Current biology, 17(3):R77–81.

Kidwell, M. G. (1983). Evolution of hybrid dysgenesis determinants in *Drosophila melanogaster*. Proceedings of the National Academy of Sciences of the United States of America, 80(6):1655–1659.

Kidwell, M. G., Kidwell, J. F., and Sved, J. A. (1977). Hybrid dysgenesis in *Drosophila melanogaster*: A syndrome of aberrant traits including mutations, sterility and male recombination. Genetics, 86(4):813–833.

Kidwell, M. G., Kimura, K., and Black, D. M. (1988). Evolution of hybrid dysgenesis potential following P element contamination in *Drosophila melanogaster*. Genetics, 119(4):815–828.

Kidwell, M. G. and Novy, J. B. (1979). Hybrid dysgenesis in *Drosophila melanogaster*: Sterility resulting from gonadal dysgenesis in the P-M system. Genetics, 92(4):1127–1140.

King, E. G., Merkes, C. M., McNeil, C. L., Hoofer, S. R., Sen, S., Broman, K. W., Long, A. D., and Macdonald, S. J. (2012). Genetic dissection of a model complex trait using the *Drosophila* Synthetic Population Resource. Genome Research, 22(8):1558–1566.

Kofler, R., Betancourt, A. J., and Schlötterer, C. (2012). Sequencing of pooled DNA samples (Pool-Seq) uncovers complex dynamics of transposable element insertions in *Drosophila melanogaster*. PLoS Genetics, 8(1):e1002487.

Kofler, R., Gómez-Sánchez, D., and Schlötterer, C. (2016). PoPoolationTE2: Comparative Population Genomics of Transposable Elements Using Pool-Seq. Molecular Biology and Evolution, 33(10):2759–2764.

Kofler, R., Hill, T., Nolte, V., Betancourt, A. J., and Schlötterer, C. (2015a). The recent invasion of natural *Drosophila simulans* populations by the P-element. Proceedings of the National Academy of Sciences of the United States of America, 112(21):6659–63.

Kofler, R., Nolte, V., and Schlötterer, C. (2015b). Tempo and Mode of Transposable Element Activity in *Drosophila*. PLoS Genetics, 11(7):1–21.

Kofler, R., Senti, K.-A., Nolte, V., Tobler, R., and Schlotterer, C. (2018). Molecular dissection of a natural transposable element invasion. Genome research, 28(6):824–835.

Kotov, A. A., Adashev, V. E., Godneeva, B. K., Ninova, M., Shatskikh, A. S., Bazylev, S. S., Aravin, A. A., and Olenina, L. V. (2019). PiRNA silencing contributes to interspecies hybrid sterility and reproductive isolation in *Drosophila melanogaster*. Nucleic Acids Research, 47(8):4255–4271.

Lack, J. B., Cardeno, C. M., Crepeau, M. W., Taylor, W., Corbett-Detig, R. B., Stevens, K. A., Langley, C. H., and Pool, J. E. (2015). The *Drosophila* genome nexus: a population genomic resource of 623 *Drosophila melanogaster* genomes, including 197 from a single ancestral range population. Genetics, 199(4):1229–1241.

Lanno, S. M., Shimshak, S. J., Peyser, R. D., Linde, S. C., and Coolon, J. D. (2019). Investigating the role of Osiris genes in *Drosophila sechellia* larval resistance to a host plant toxin. Ecology and Evolution, 9(4):1922–1933.

Le Rouzic, A. and Capy, P. (2005). The first steps of transposable elements invasion: Parasitic strategy vs. genetic drift. Genetics, 169(2):1033–1043.

Le Thomas, A., Rogers, A. K., Webster, A., Marinov, G. K., Liao, S. E., Perkins, E. M., Hur, J. K., Aravin, A. A., and Tóth, K. F. (2013). Piwi induces piRNA-guided transcriptional silencing and establishment of a repressive chromatin state. Genes Dev., 27(4):390–399.

Lemeunier, F., Ashburner, M., and Thoday, J. M. (1976). Relationships within the *melanogaster* species subgroup of the genus *Drosophila (Sophophora*) - II. Phylogenetic relationships between six species based upon polytene chromosome banding sequences. Proceedings of the Royal Society of London. Series B. Biological Sciences, 193(1112):275–294.

Lerat, E., Burlet, N., Biémont, C., and Vieira, C. (2011). Comparative analysis of transposable elements in the *melanogaster* subgroup sequenced genomes. Gene, 473(2):100–109.

Li, H. and Durbin, R. (2009). Fast and accurate short read alignment with Burrows–Wheeler transform. Bioinformatics, 25(14):1754–1760.

Li, H. and Durbin, R. (2010). Fast and accurate long-read alignment with Burrows-Wheeler transform. Bioinformatics, 26(5):589–95.

Li, H., Handsaker, B., Wysoker, A., Fennell, T., Ruan, J., Homer, N., Marth, G., Abecasis, G., and Durbin, R. (2009). The Sequence Alignment/Map format and SAMtools. Bioinformatics, 25(16):2078–9.

Lindsley, D. H. and Grell, E. H. (1968). Genetic variations of Drosophila melanogaster. Carnegie Institute of Washington Publication.

Lohe, A. R., Moriyama, E. N., Lidholm, D. A., and Hartl, D. L. (1995). Horizontal transmission, vertical inactivation, and stochastic loss of mariner-like transposable elements. Molecular Biology and Evolution, 12(1):62–72.

Loreto, E. L., Carareto, C. M. A., and Capy, P. (2008). Revisiting horizontal transfer of transposable elements in *Drosophila*. Heredity, 100(6):545–554.

Machado, H. E., Bergland, A. O., Taylor, R., Tilk, S., Behrman, E., Dyer, K., Fabian, D. K., Flatt, T., González, J., Karasov, T. L., et al. (2019). Broad geographic sampling reveals predictable, pervasive, and strong seasonal adaptation in *Drosophila*. bioRxiv, page 337543.

Mackay, T. F., Richards, S., Stone, E. A., Barbadilla, A., Ayroles, J. F., Zhu, D., Casillas, S., Han, Y., Magwire, M. M., Cridland, J. M., et al. (2012). The *Drosophila melanogaster* genetic reference panel. Nature, 482(7384):173–178.

Malone, C. D., Brennecke, J., Dus, M., Stark, A., McCombie, W. R., Sachidanandam, R., and Hannon, G. J. (2009). Specialized piRNA Pathways Act in Germline and Somatic Tissues of the *Drosophila* Ovary. Cell, 137(3):522–535.

Maniatis, T., Fritsch, E. F., Sambrook, J., et al. (1982). Molecular cloning: a laboratory manual, volume 545. Cold spring harbor laboratory Cold Spring Harbor, NY.

Marí-Ordóñez, A., Marchais, A., Etcheverry, M., Martin, A., Colot, V., and Voinnet, O. (2013). Reconstructing *de novo* silencing of an active plant retrotransposon. Nature genetics, 45(9):1029–1039.

Marsano, R. M., Moschetti, R., Caggese, C., Lanave, C., Barsanti, P., and Caizzi, R. (2000). The complete Tirant transposable element in *Drososphila melanogaster* shows a structural relationship with retrovirus-like retrotransposons. Gene, 247(1-2):87–95.

Martin, M. (2011). Cutadapt removes adapter sequences from high-throughput sequencing reads. EMBnet.journal, 17(1):10–12.

Maruyama, K. and Hartl, D. L. (1991). Evidence for interspecific transfer of the transposable element mariner between *Drosophila* and *Zaprionus*. Journal of Molecular Evolution, 33(6):514–524.

Meany, M. K., Conner, W. R., Richter, S. V., Bailey, J. A., Turelli, M., and Cooper, B. S. (2019). Loss of cytoplasmic incompatibility and minimal fecundity effects explain relatively low Wolbachia frequencies in *Drosophila mauritiana*. Evolution, 73(6):1278–1295.

Melvin, R. G., Lamichane, N., Havula, E., Kokki, K., Soeder, C., Jones, C. D., and Hietakangas, V. (2018). Natural variation in sugar tolerance associates with changes in signaling and mitochondrial ribosome biogenesis. eLife, 7:e40841.

Miller, D. E., Staber, C., Zeitlinger, J., and Hawley, R. S. (2018). Highly Contiguous Genome Assemblies of 15 *Drosophila* Species Generated Using Nanopore Sequencing. G3; GenesȔGenomesȔGenetics, 8(10):3131–3141.

Mizrokhi, L. J. and Mazo, A. M. (1990). Evidence for horizontal transmission of the mobile element jockey between distant *Drosophila* species. Proceedings of the National Academy of Sciences, 87(23):9216–9220.

Moltó, M. D., Paricio, N., López-Preciado, M. a., Semeshin, V. F., and Martínez-Sebastián, M. J. (1996). Tirant: a new retrotransposon-like element in *Drosophila melanogaster*. Journal of molecular evolution, 42(3):369–75.

Moon, S., Cassani, M., Lin, Y. A., Wang, L., Dou, K., and Zhang, Z. Z. (2018). A Robust Transposon-Endogenizing Response from Germline Stem Cells. Developmental Cell, 47(5):660–671.e3.

Mugnier, N., Gueguen, L., Vieira, C., and Biémont, C. (2008). The heterochromatic copies of the LTR retrotransposons as a record of the genomic events that have shaped the *Drosophila melanogaster* genome. Gene, 411(1-2):87–93.

Orgel, L. E. and Crick, F. H. (1980). Selfish DNA: the ultimate parasite. Nature, 284(5757):604–607.

Pardue, M. L. and DeBaryshe, P. G. (2011). Retrotransposons that maintain chromosome ends. Proceedings of the National Academy of Sciences of the United States of America, 108(51):20317–20324.

Parsons, P. and Stanley, S. (1981). Special ecological studies-domesticated and widespread species. In Ashburner, M., Carson, L., and Thompson, J. J., editors, The genetics and biology of Drosophila, volume 3c, pages 349–393. Academic Press, Oxford.

Peccoud, J., Loiseau, V., Cordaux, and Gilbert, C. (2017). Massive horizontal transfer of transposable elements in insects. PNAS, 114(18):4721–4726.

Periquet, G., Hamelin, M. H., Bigot, Y., and Lepissier, A. (1989). Geographical and historical patterns of distribution of hobo elements in *Drosophila melanogaster* populations. Journal of Evolutionary Biology, 2(3):223–229.

Periquet, G., Lemeunier, F., Bigot, Y., Hamelin, M. H., Bazin, C., Ladevèze, V., Eeken, J., Galindo, M. I., Pascual, L., and Boussy, I. (1994). The evolutionary genetics of the hobo transposable element in the *Drosophila melanogaster* complex. Genetica, 93(1-3):79–90.

Petrov, D. A., Fiston-Lavier, A.-S., Lipatov, M., Lenkov, K., and González, J. (2011). Population genomics of transposable elements in *Drosophila melanogaster*. Molecular biology and evolution, 28(5):1633–44.

Post, C., Clark, J. P., Sytnikova, Y. A., Chirn, G.-w., and Lau, N. C. (2014). The capacity of target silencing by *Drosophila* PIWI and piRNAs The capacity of target silencing by *Drosophila* PIWI and piRNAs. RNA, 20(12):1977–1986.

Quesneville, H., Bergman, C. M., Andrieu, O., Autard, D., Nouaud, D., Ashburner, M., and Anxolabéhère, D. (2005). Combined evidence annotation of transposable elements in genome sequences. PLoS computational biology, 1(2):166–175.

Quinlan, A. R. and Hall, I. M. (2010). BEDTools: a flexible suite of utilities for comparing genomic features. Bioinformatics, 26(6):841–842.

R Core Team (2012). R: A Language and Environment for Statistical Computing. R Foundation for Statistical Computing, Vienna, Austria. ISBN 3-900051-07-0.

Rahman, R., Chirn, G. W., Kanodia, A., Sytnikova, Y. A., Brembs, B., Bergman, C. M., and Lau, N. C. (2015). Unique transposon landscapes are pervasive across *Drosophila melanogaster* genomes. Nucleic Acids Research, 43(22):10655–10672.

Riddle, N. C., Minoda, A., Kharchenko, P. V., Alekseyenko, A. A., Schwartz, Y. B., Tolstorukov, M. Y., Gorchakov, A. A., Jaffe, J. D., Kennedy, C., Linder-Basso, D., et al. (2011). Plasticity in patterns of histone modifications and chromosomal proteins in *Drosophila* heterochromatin. Genome Research, 21(2):147–163.

Rogers, R. L., Cridland, J. M., Shao, L., Hu, T. T., Andolfatto, P., and Thornton, K. R. (2014). Landscape of standing variation for tandem duplications in *Drosophila yakuba* and *Drosophila simulans*. Molecular biology and evolution, 31(7):1750–1766.

Ruebenbauer, A., Schlyter, F., Hansson, B. S., Löfstedt, C., and Larsson, M. C. (2008). Genetic Variability and Robustness of Host Odor Preference in *Drosophila melanogaster*. Current Biology, 18(18):1438–1443.

Saint-Leandre, B. and Levine, M. T. (2020). The Telomere Paradox: Stable Genome Preservation with Rapidly Evolving Proteins. Trends in Genetics, 36(4):232–242.

Sánchez-Gracia, A., Maside, X., and Charlesworth, B. (2005). High rate of horizontal transfer of transposable elements in *Drosophila*. Trends in genetics, 21(4):200–203.

Schrider, D., Ayroles, J., Matute, D., and Kern, A. (2018). Supervised machine learning reveals introgressed loci in the genomes of *Drosophila simulans* and *D. sechellia*. PLOS Genetics, 14(4):170670.

Serrato-Capuchina, A., Wang, J., Earley, E., Peede, D., Isbell, K., and Matute, D. (2020). Paternally inherited P -element copy number affects the magnitude of hybrid dysgenesis in *Drosophila simulans* and *D. melanogaster*. Genome Biology and Evolution, page evaa084.

Sienski, G., Dönertas, D., and Brennecke, J. (2012). Transcriptional silencing of transposons by Piwi and Maelstrom and its impact on chromatin state and gene expression. Cell, 151(5):964–980.

Silva, J. C., Loreto, E. L., and Clark, J. B. (2004). Factors that affect the horizontal transfer of transposable elements. Curr Issues Mol Biol, 6(1):57–72.

Simmons, G. M. (1992). Horizontal transfer of hobo transposable elements within the *Drosophila melanogaster* species complex: evidence from DNA sequencing. Molecular Biology and Evolution, 9(6):1050–1060.

Smit, A. F. A., Hubley, R., and Green, P. (2013-2015). RepeatMasker Open-4.0.

Sniegowski, P. D. and Charlesworth, B. (1994). Transposable element numbers in cosmopolitan inversions from a natural population of *Drosophila melanogaster*. Genetics, 137(3):815–827.

Song, S. U., Kurkulos, M., Boeke, J. D., and Corces, V. G. (1997). Infection of the germ line by retroviral particles produced in the follicle cells: a possible mechanism for the mobilization of the gypsy retroelement of *Drosophila*. Development, 124(14):2789–2798.

Srivastav, S. P., Rahman, R., Ma, Q., Pierre, J., Bandyopadhyay, S., and Lau, N. C. (2019). Har-P, a short P-element variant, weaponizes P-transposase to severely impair *Drosophila* development. eLife, 26(6):e49948.

Stewart, N. B. and Rogers, R. L. (2019). Chromosomal rearrangements as a source of new gene formation in *Drosophila yakuba*. PLOS Genetics, 15(9):e1008314.

Terzian, C., Ferraz, C., Demaille, J., and Bucheton, A. (2000). Evolution of the Gypsy Endogenous Retrovirus in the *Drosophila melanogaster* Subgroup. Molecular Biology and Evolution, 17(6):908–914.

Terzian, C., Pélisson, A., and Bucheton, A. (2001). Evolution and phylogeny of insect endogenous retroviruses. BMC Evolutionary Biology, 1(3).

Thurmond, J., Goodman, J. L., Strelets, V. B., Attrill, H., Gramates, L. S., Marygold, S. J., Matthews, B. B., Millburn, G., Antonazzo, G., Trovisco, V., Kaufman, T. C., Calvi, B. R., Perrimon, N., Gelbart, S. R., Agapite, J., Broll, K., Crosby, L., Dos Santos, G., Emmert, D., Falls, K., Jenkins, V., Sutherland, C., Tabone, C., Zhou, P., Zytkovicz, M., Brown, N., Garapati, P., Holmes, A., Larkin, A., Pilgrim, C., Urbano, P., Czoch, B., Cripps, R., and Baker, P. (2019). FlyBase 2.0: The next generation. Nucleic Acids Research, 47(D1):D759–D765.

Turissini, D. A., Liu, G., David, J. R., and Matute, D. R. (2015). The evolution of reproductive isolation in the *Drosophila yakuba* complex of species. Journal of Evolutionary Biology, 28(3):557–575.

Vieira, C., Lepetit, D., Dumont, S., and Biémont, C. (1999). Wake up of transposable elements following *Drosophila simulans* worldwide colonization. Molecular biology and evolution, 16(9):1251–1255.

Viggiano, L., Caggese, C., Barsanti, P., and Caizzi, R. (1997). Cloning and characterization of a copy of Tirant transposable element in *Drosophila melanogaster*. Gene, 197(1-2):29–35.

Wang, L., Dou, K., Moon, S., Tan, F. J., Zhang, Z. Z. Z., Wang, L., Dou, K., Moon, S., Tan, F. J., and Zhang, Z. Z. Z. (2018). Hijacking Oogenesis Enables Massive Propagation Article Hijacking Oogenesis Enables Massive Propagation of LINE and Retroviral Transposons. Cell, 174(5):1082–1094.

Weilguny, L. and Kofler, R. (2019). DeviaTE: Assembly-free analysis and visualization of mobile genetic element composition. Molecular Ecology Resources, 19(5):1346–1354.

Wicker, T., Sabot, F., Hua-Van, A., Bennetzen, J. L., Capy, P., Chalhoub, B., Flavell, A., Leroy, P., Morgante, M., Panaud, O., et al. (2007). A unified classification system for eukaryotic transposable elements. Nature Reviews Genetics, 8(12):973–982.

Wierzbicki, F., Schwarz, F., Cannalonga, O., and Kofler, R. (2020). Generating high quality assemblies for genomic analysis of transposable elements. bioRxiv.

Yang, P., Wang, Y., and Macfarlan, T. S. (2017). The Role of KRAB-ZFPs in Transposable Element Repression and Mammalian Evolution. Trends Genet., 33(11):871–881.

Yannopoulos, G., Stamatis, N., Monastirioti, M., Hatzopoulos, P., and Louis, C. (1987). hobo is responsible for the induction of hybrid dysgenesis by strains of *Drosophila melanogaster* bearing the male recombination factor 23.5MRF. Cell, 49(4):487–495.

