## supplementary material for "Tirant stealthily invaded natural *Drosophila melanogaster* populations during the last century"

June 10, 2020

### Supplementary figures

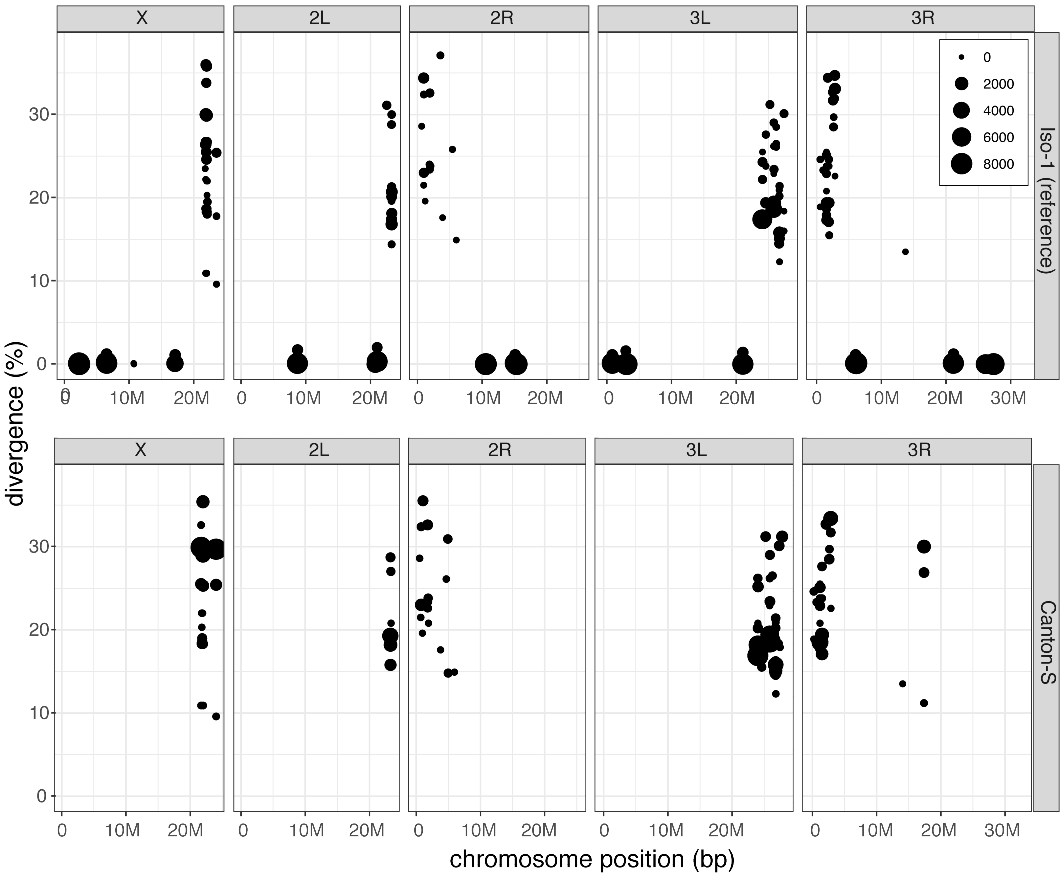


Figure 1: Canonical Tirant insertions are present in Iso-1 but not Canton-S. The Canton-S assembly was generated by Chakraborty et al. (2019) with PacBio reads (the Canton-S assembly shown in the main manuscript was generated by Wierzbicki et al. (2020) with ONT reads). For each Tirant insertion we show the position in the assembly, the length (size of dot), and the similarity to the consensus sequence (divergence).

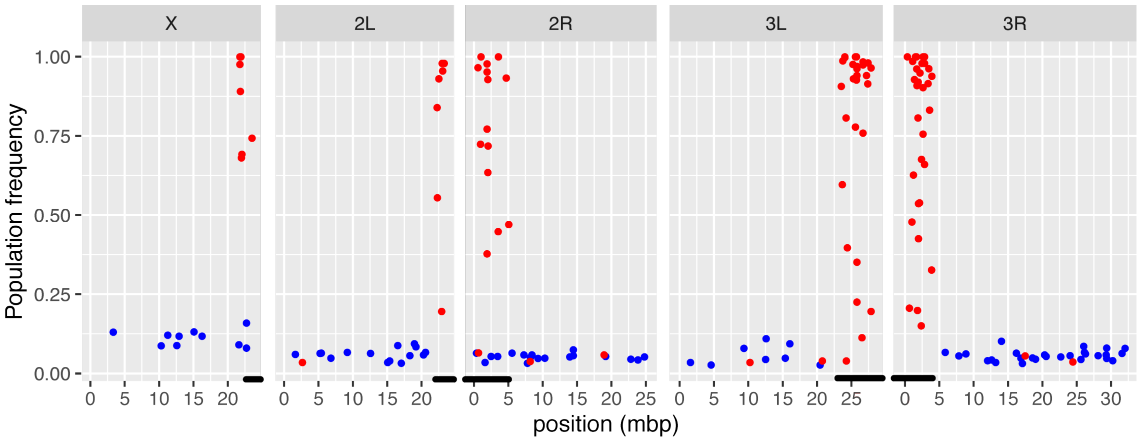


Figure 2: Position and population frequency of canonical (blue) and degraded (red) Tirant insertions in a population from France (Viltain) (Kapun et al., 2018). Canonical Tirant insertions are mostly euchromatic and segregating at a low population frequency whereas degraded insertions are mostly heterochromatic and segregating at high frequency. Black bars indicate (peri)centric heterochromatin (Riddle et al., 2011; Hoskins et al., 2015).

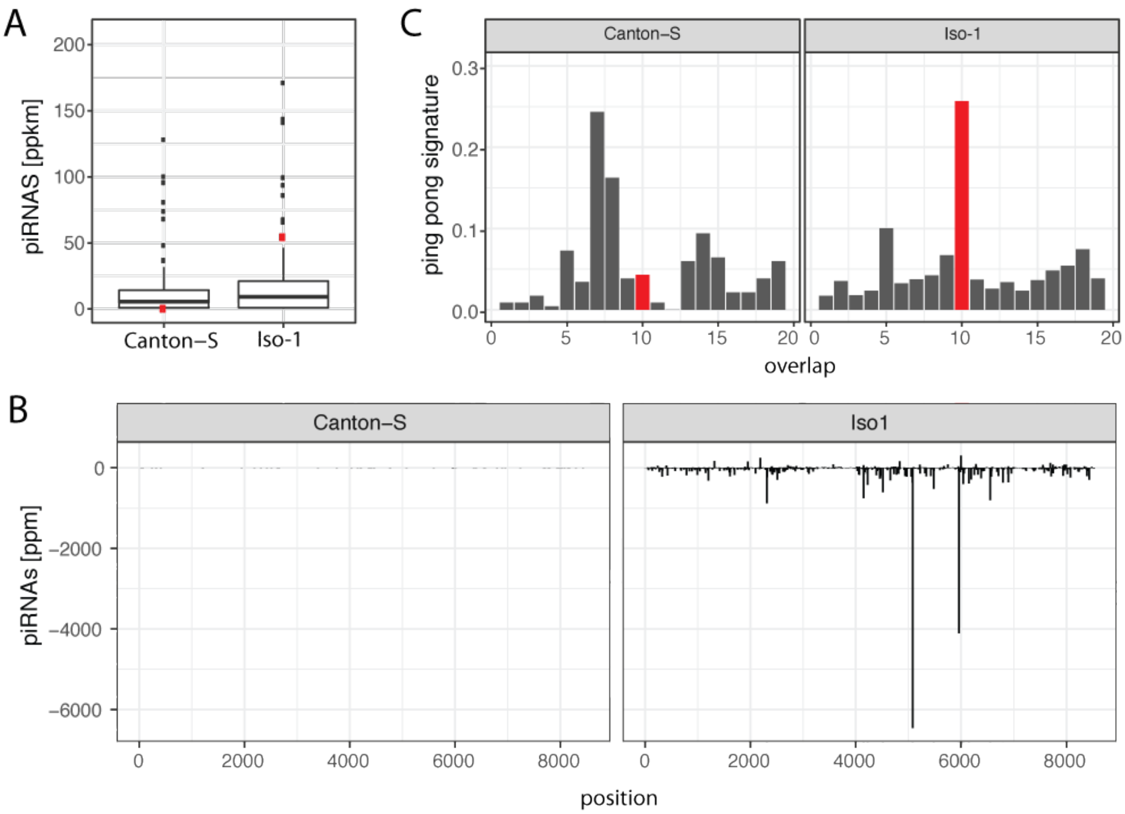


Figure 3: Tirant piRNAs in Iso-1 and Canton-S. A) Abundance of Tirant piRNAs (red) compared to piRNAs complementary to the other TEs of *D. melanogaster*. B) Abundance of sense (positive y-axis) and antisense (negative y-axis) piRNAs along the sequence of Tirant. C) Ping-pong signature for Iso-1 and Canton-S. A pronounced peak at position 10 (red) suggest secondary amplification of piRNAs by the ping-pong cycle.

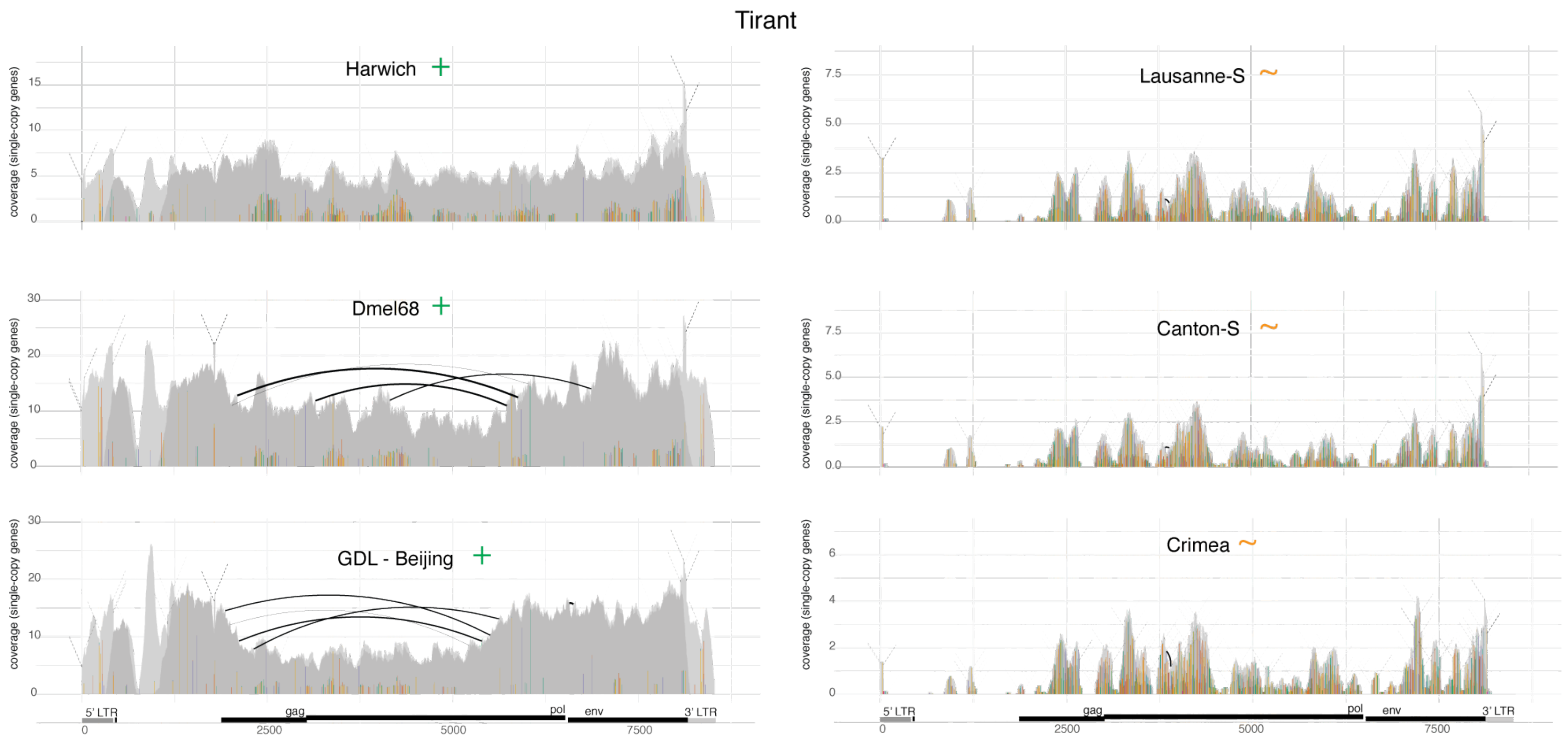


Figure 4: DeviaTE plots for three strains having non-degraded (i.e. canonical) Tirant sequences (+) and three strains solely having degraded Tirant sequences (~). Such plots were used for classifying the Tirant content of the different *D. melanogaster* strains (see supplementary table 1).

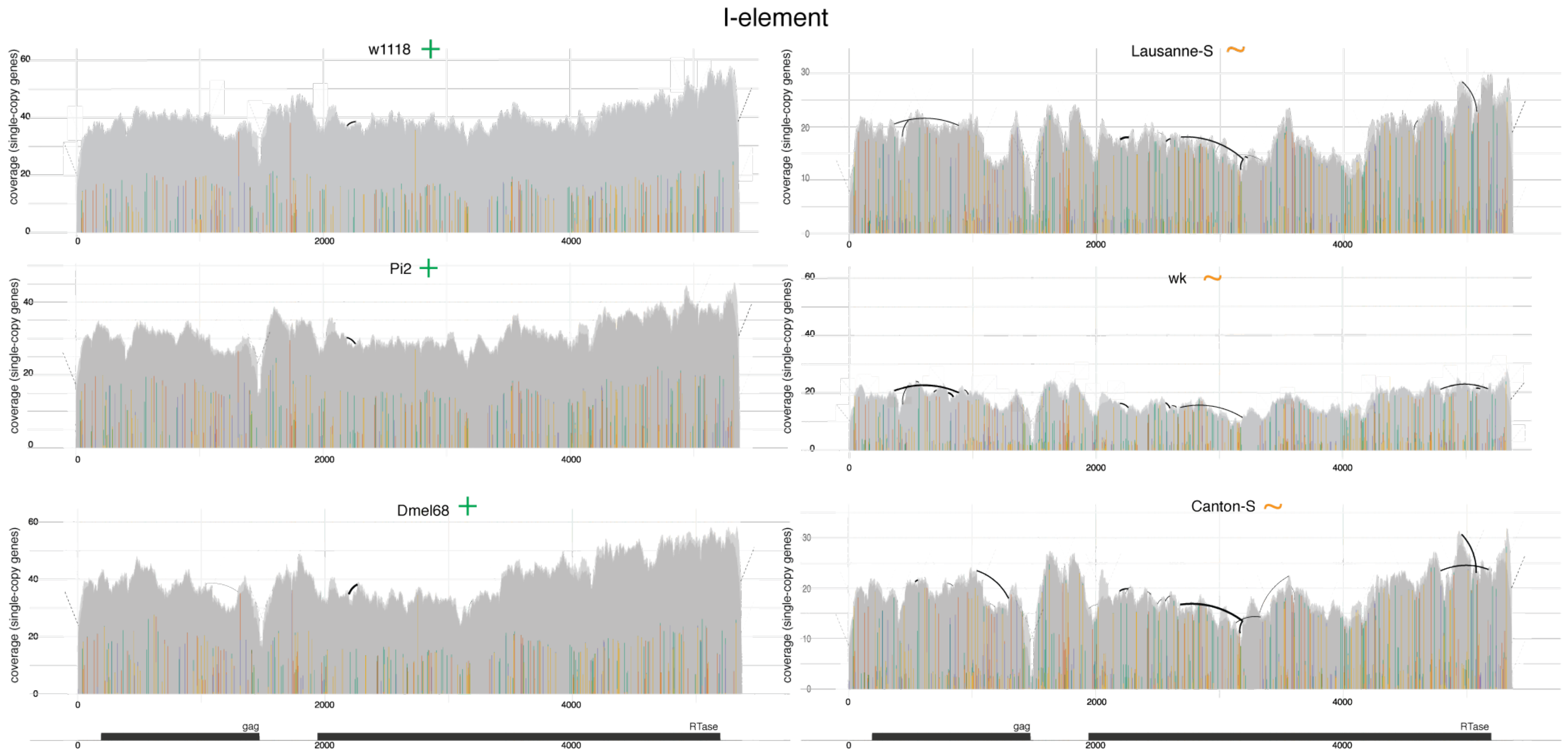


Figure 5: DeviaTE plots for three strains having non-degraded I-element sequences (+) and three strains solely having degraded I-element sequences (~). Such plots were used for classifying the I-element content of the different *D. melanogaster* strains (see supplementary table 1).

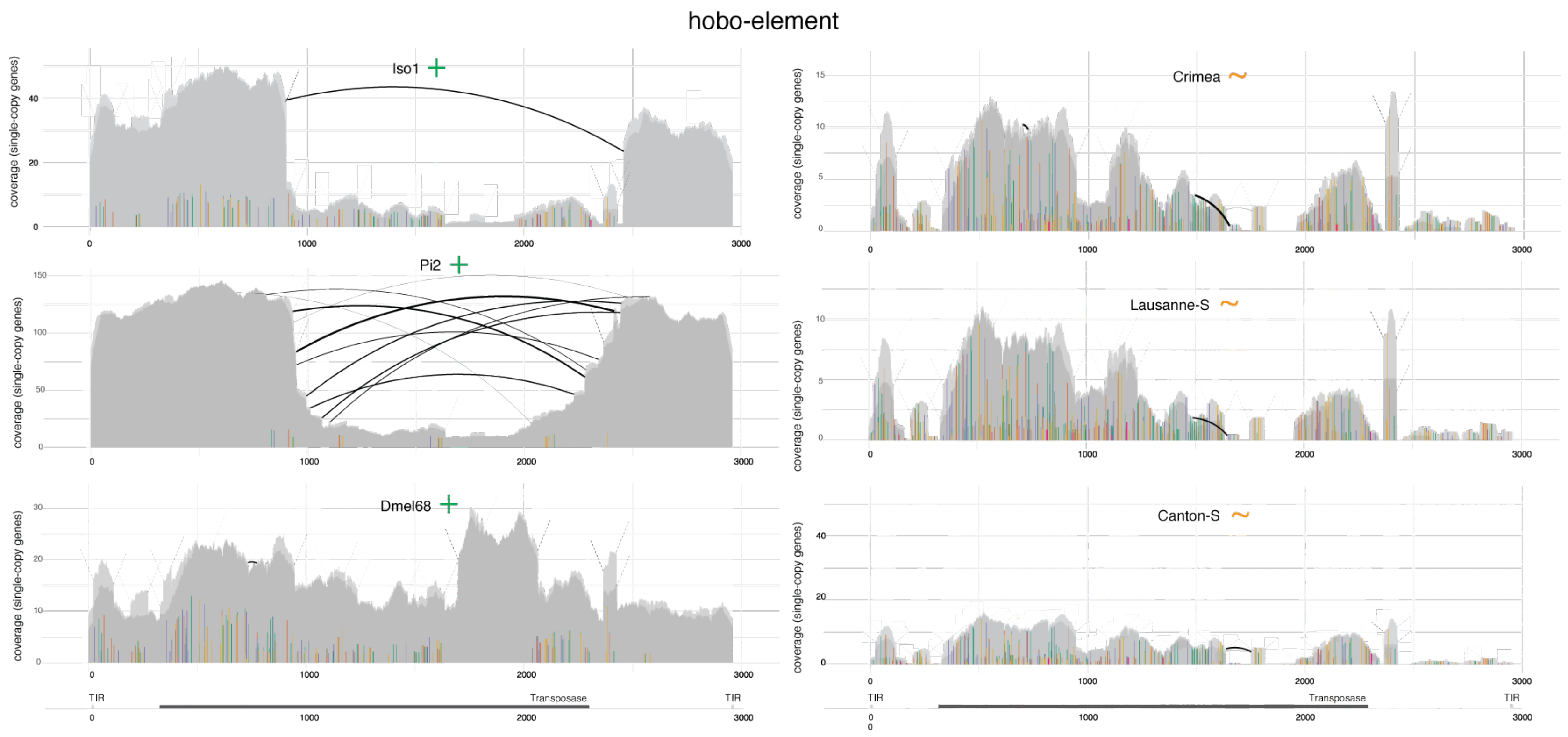


Figure 6: DeviaTE plots for three strains having non-degraded hobo sequences (+) and three strains solely having degraded hobo sequences (~). Such plots were used for classifying the hobo content of the different *D. melanogaster* strains (see supplementary table 1).

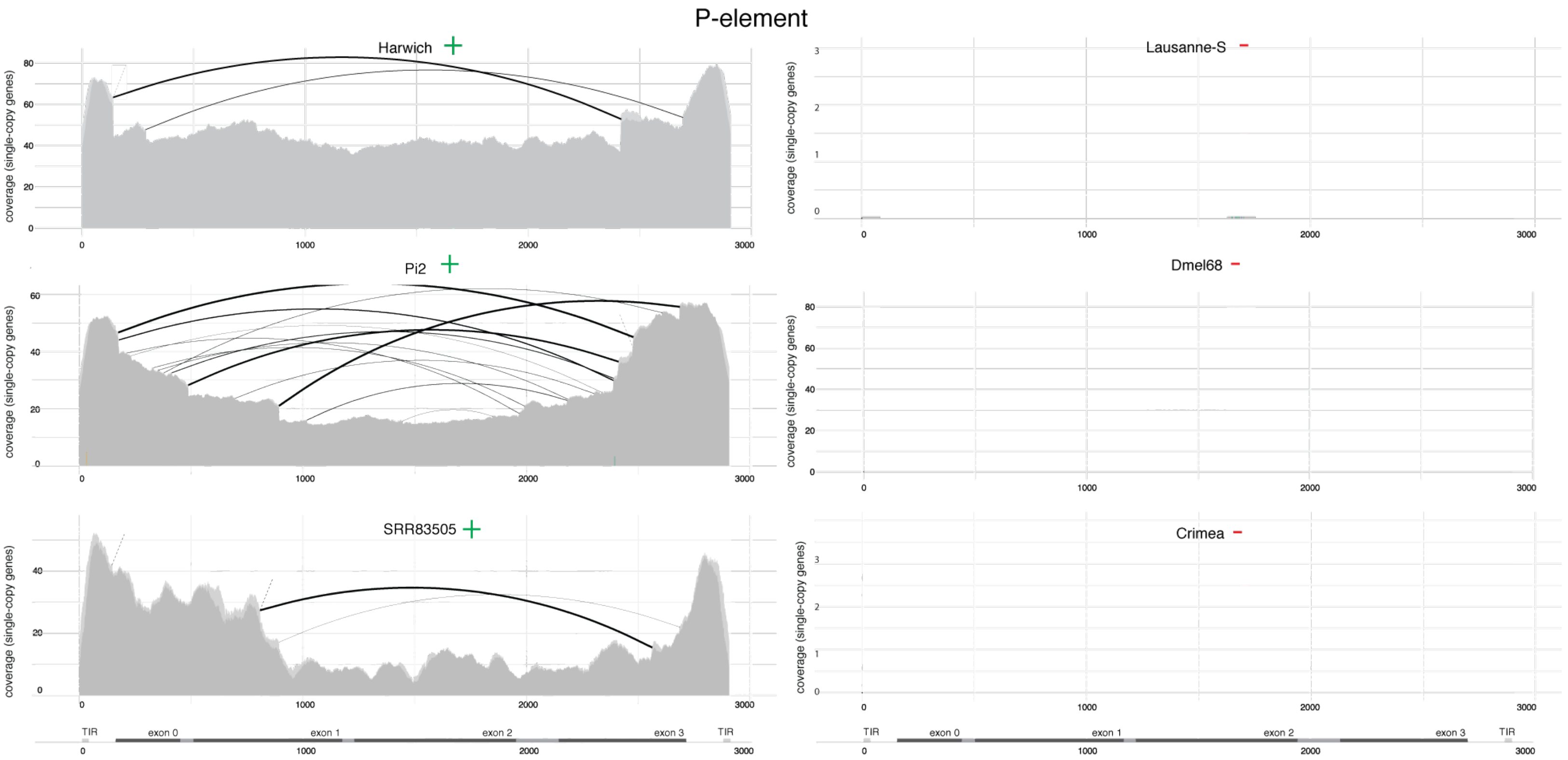


Figure 7: DeviaTE plots for three strains having P-element sequences (+) and three strains not having P-element sequences (−). Such plots were used for classifying the P-element content of the different *D. melanogaster* strains (see supplementary table 1).

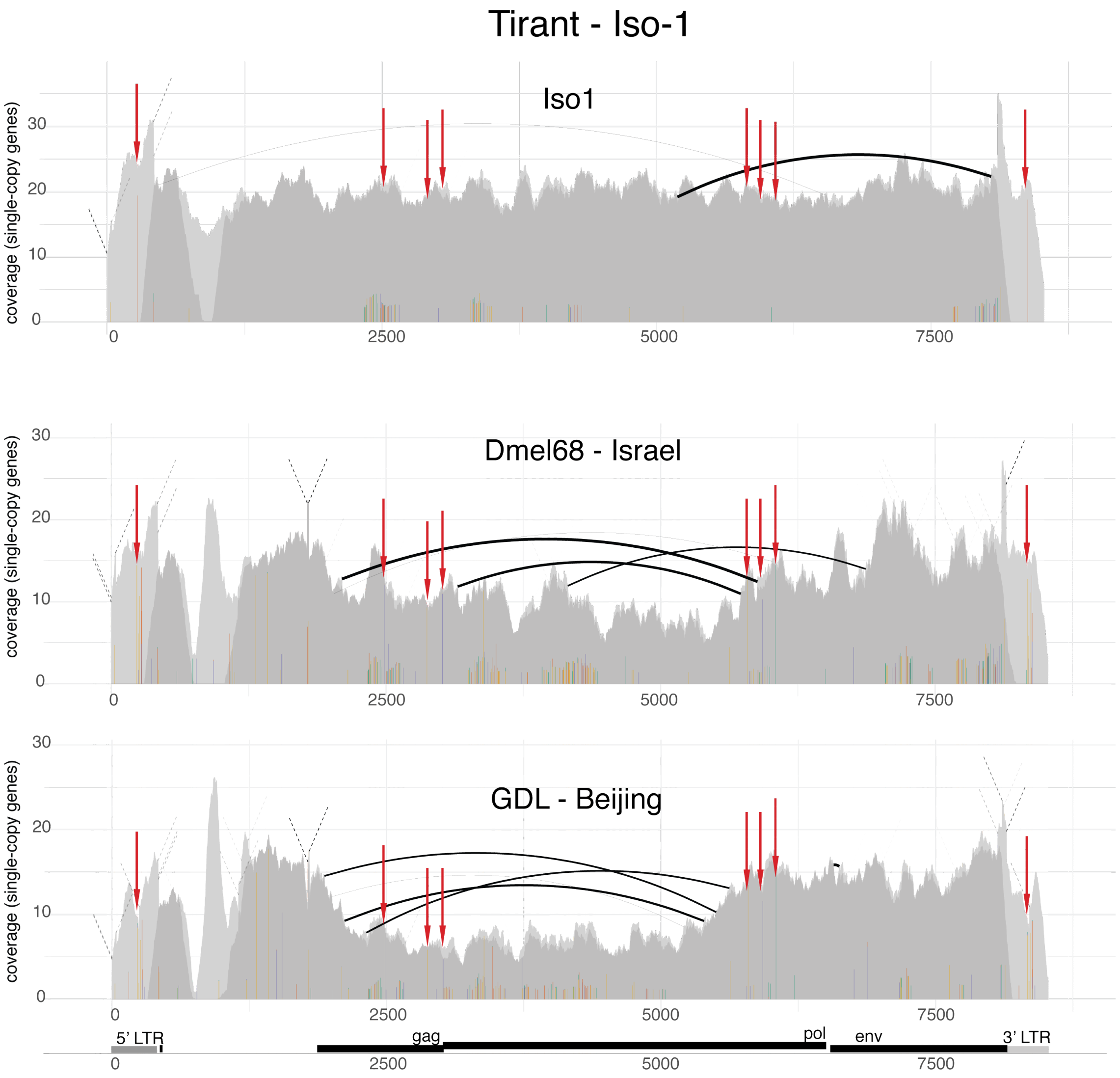


Figure 8: Abundance and diversity of Tirant in the reference strain Iso-1 and two strains collected from natural *D. melanogaster* populations (Dmel68 and a GDL line from Beijing). Eight SNPs found in natural populations but not in Iso-1 are marked by red arrows.

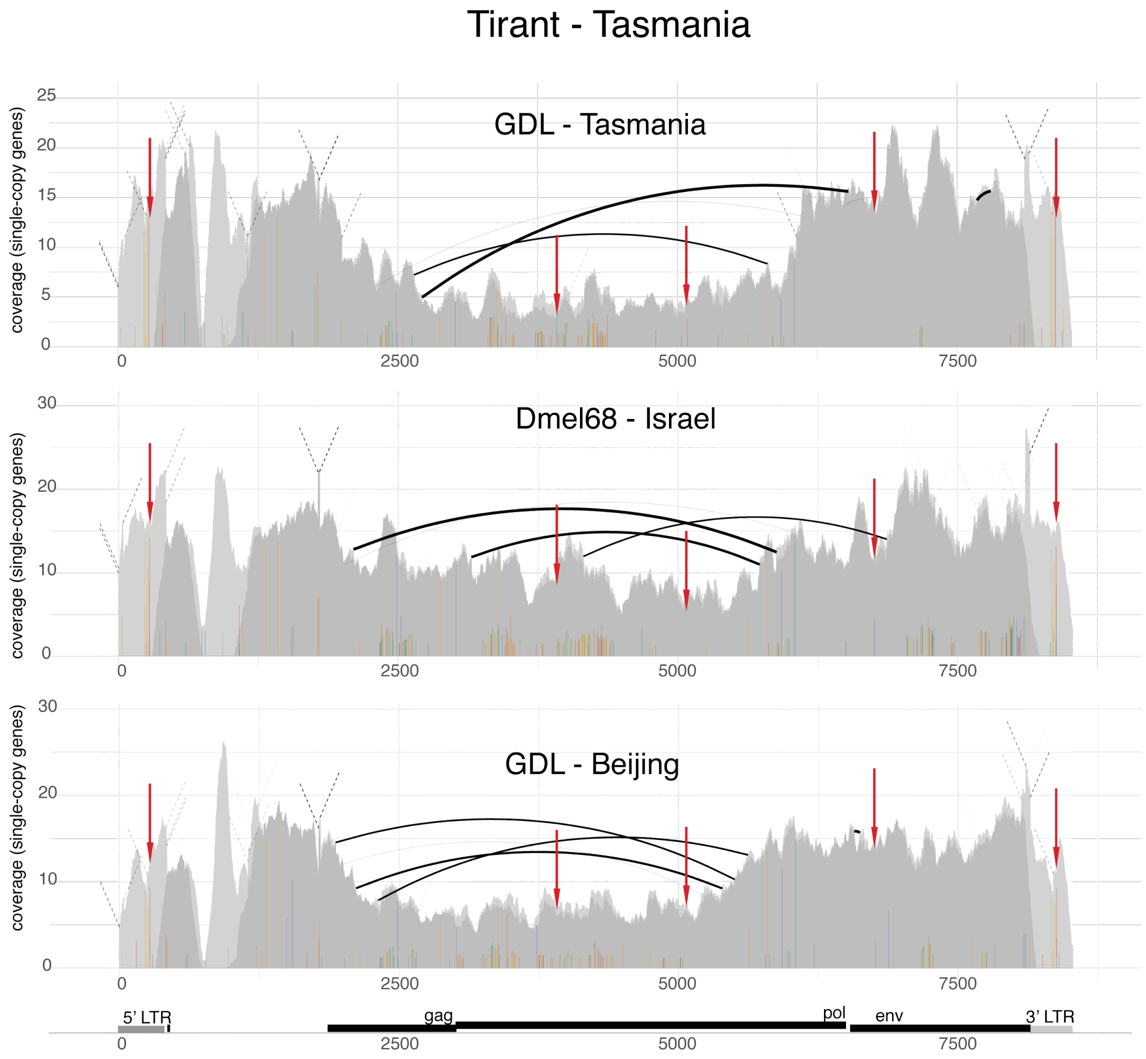


Figure 9: Abundance and diversity of Tirant sequences in a natural population from Tasmania and from other geographic locations. Five SNPs, marked by red arrows, have notably different allele frequencies between populations from Tasmania and the other geographic locations.

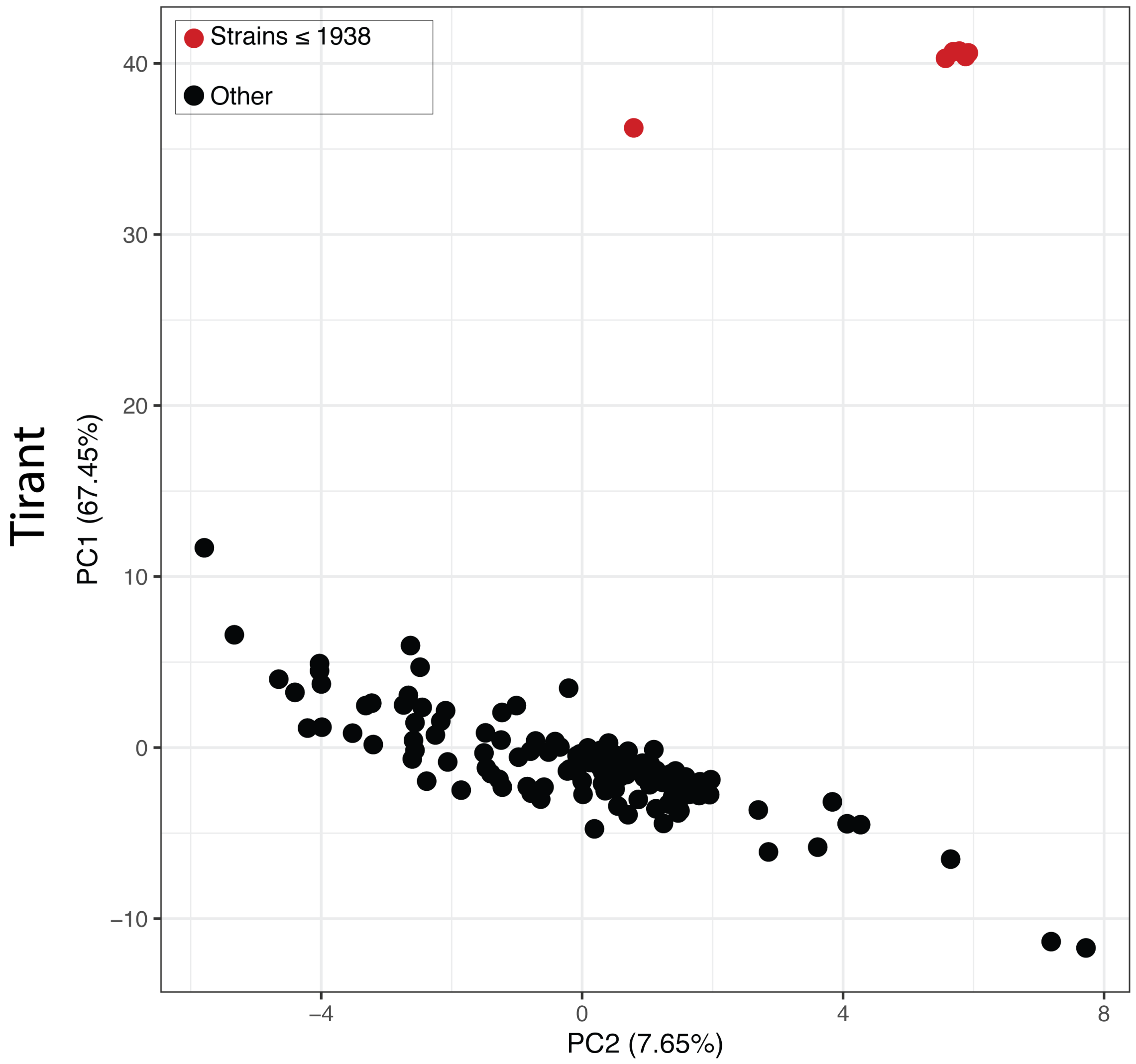


Figure 10: PCA based on the allele frequencies of SNPs in Tirant for different *D. melanogaster* strains and population samples. Strains sampled before or at 1938 form a separate cluster (due to the absence of canonical Tirant insrtions). In addition to the strains shown in the manuscript (fig. 3) we used DGRP, DrosEU and Dros-RTEC lines well as lines sampled by Bergland et al. (2014) and Lack et al. (2015) (see supplementary table 1 for details).

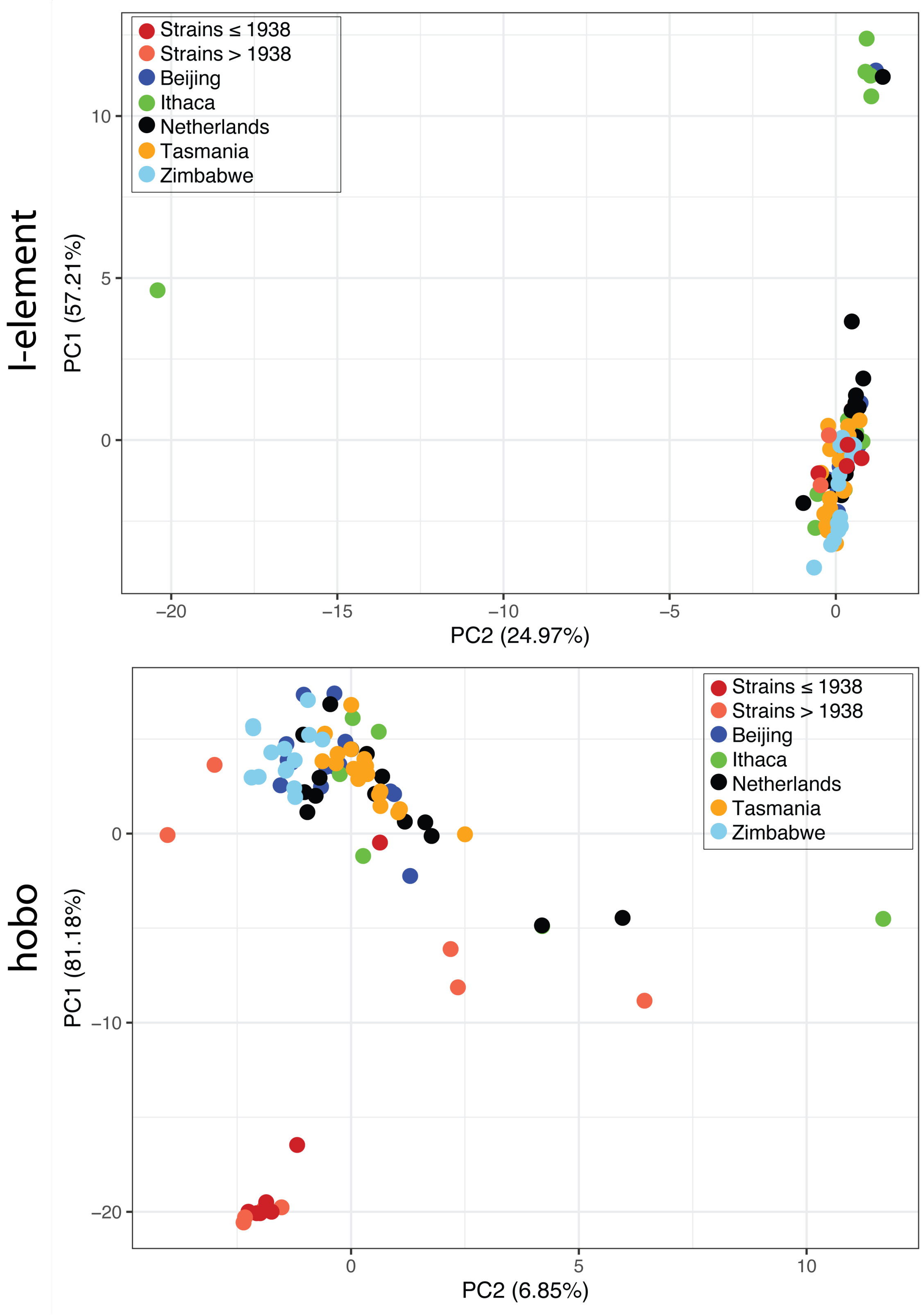


Figure 11: PCA based on the allele frequencies of SNPs in the I-element and hobo. Tasmanian populations cluster with strains from other geographic regions for both TEs.

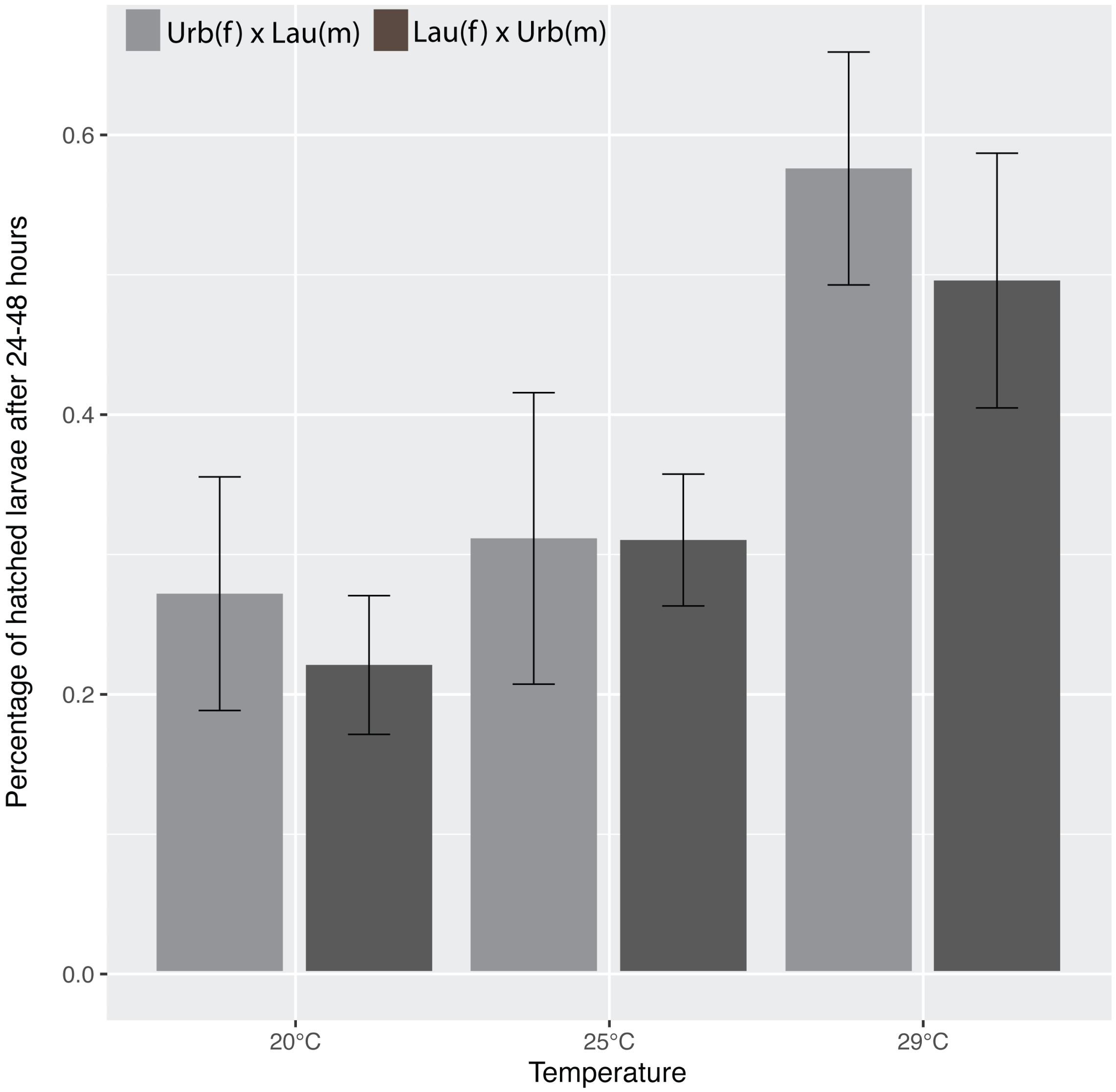


Figure 12: Fraction of hatched F2 eggs for reciprocal crosses between a strain having recent Tirant insertions (Urb: Urbana-S) and a strain not having recent Tirant insertions (Lau: Laussane-S). Crosses were performed at three temperatures and three replicates were used for each cross. We did not detect significant differences in the abundance of hatched F2 eggs between the reciprocal crosses (Wilcoxon rank sum test; $p_{20} = 0.4$, $p_{25} = 0.7$, $p_{29} = 0.4$). Differences in absolute number of hatched eggs between the temperatures are caused by an increased hatching rate at higher temperatures; m males, f females. .

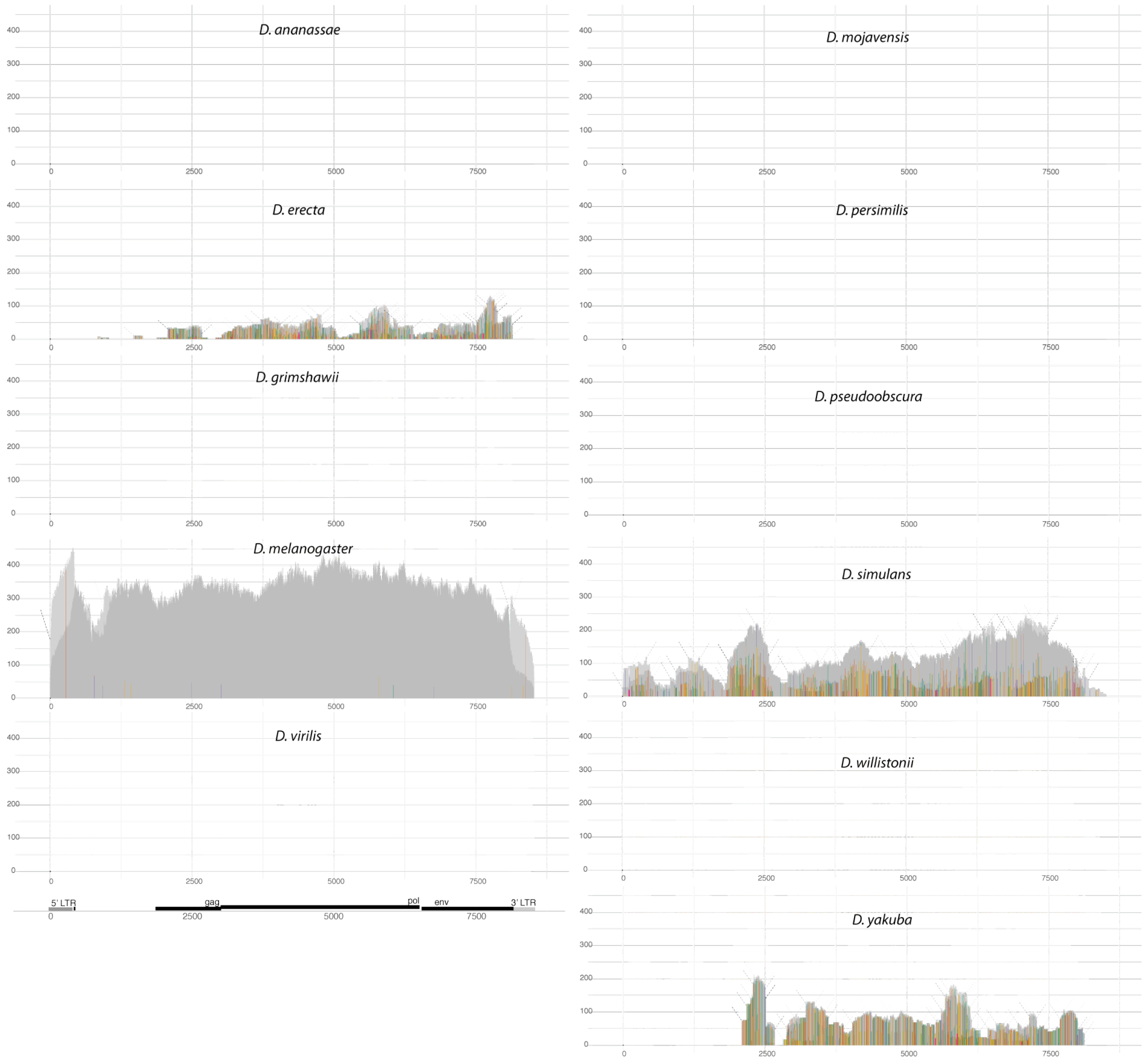


Figure 13: Abundance and diversity of Tirant sequences in 11 Drosophila species (Drosophila 12 Genomes Consortium, 2007). Tirant sequences can solely be found in the *Drosophila melanogaster* species subgroup.

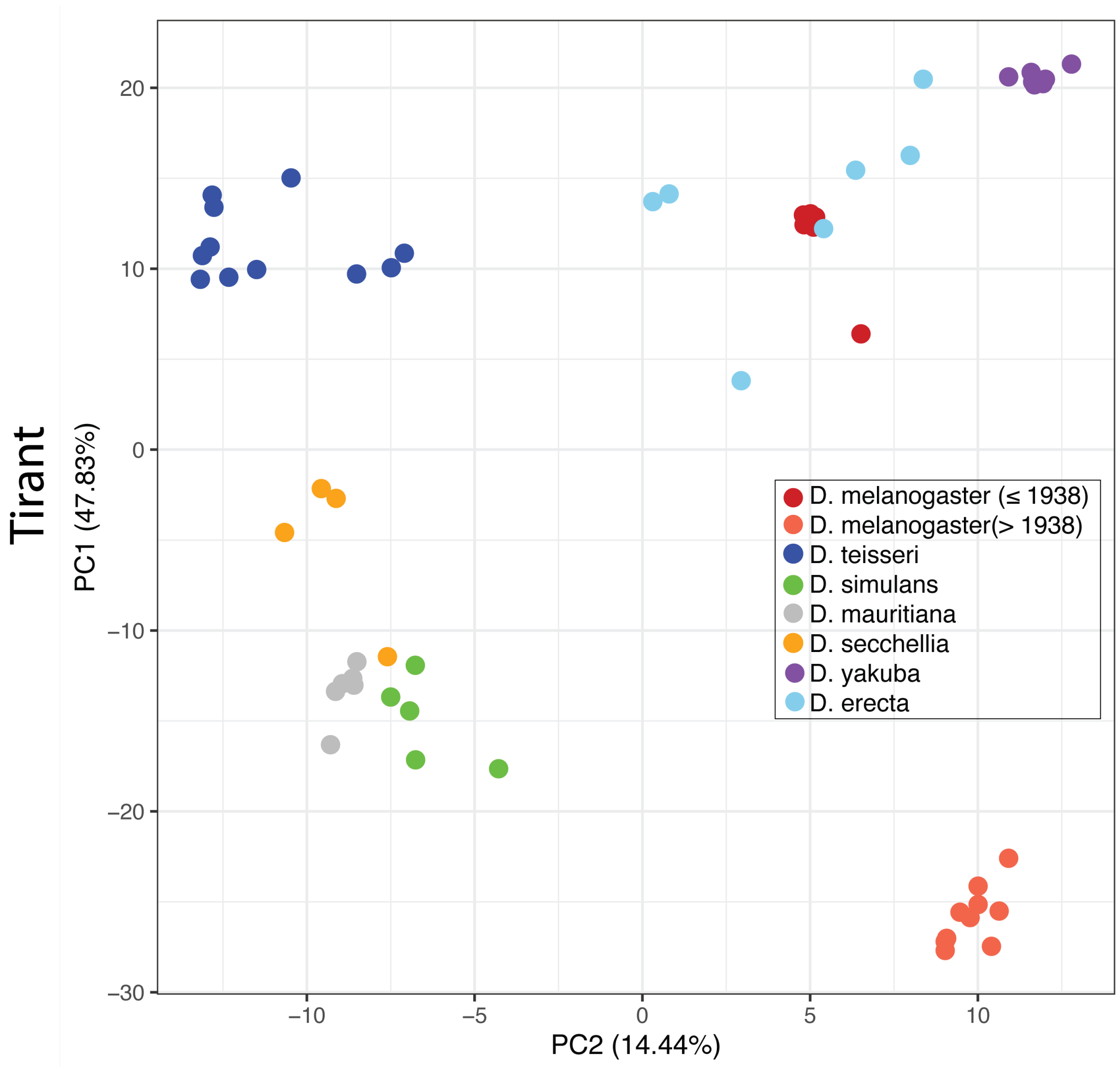


Figure 14: PCA based on the allele frequencies of SNPs in Tirant. Data are shown for several lines of different species from the *Drosophila melanogaster* species subgroup. Note that old lab strains of *D. melanogaster* cluster with *D. erecta* while more recently collected strain are closest to *D. simulans*.

### Supplementary tables

Table 1: Overview of the abundance of Tirant, I-element, hobo, and P-element sequences in different *D. melanogaster* strains. Strains are ordered by their estimated collection date. For each family and strain, we classified the TE content into three distinct categories: 'red' absence of any TE sequence, 'yellow' solely degraded TE sequences are present, 'green' non-degraded sequences, with a high similarity to the consensus sequence are present. Numbers in brackets represent the average coverage normalized to single-copy genes (≈ TE copy numbers per haploid genome). Strains sequenced in this work are marked by a star (*). † latest possible collection date was inferred from death of C. Bridges (1938), who collected the strain (Lindsley and Grell, 1968). coll. date collection date, FlyBase `https://flybase.org/`, NDSSC `https://www.drosophilaspecies.com/`

| strain | coll. date | Tirant | I-ele. | hobo | P-ele. | location | source |
|---|---|---|---|---|---|---|---|
| Oregon-R | 1925 | ~ (0.6) | ~ (19.9) | ~ (5.8) | − (0) | Oregon, USA | Lindsley and Grell 1968 |
| Canton-S | 1935 | ~ (0.9) | ~ (19.5) | ~ (5.9) | − (0) | Ohio, USA | Anxolabéhère *et al.* 1988 |
| Samarkand | 1936 | ~ (0.7) | ~ (17.6) | ~ (4.3) | − (0) | Samarkand, Uzbekistan | Lindsley and Grell 1968 |
| Crimea* | 1936 | ~ (0.5) | ~ (18.5) | ~ (4.5) | − (0) | Crimea, Eastern Europe | Anxolabéhère *et al.* 1988 |
| Lausanne-S* | 1938 | ~ (0.5) | ~ (18.8) | ~ (3.7) | − (0) | Wisconsin, USA | Lindsley and Grell 1968 |
| Swedish-C* | <1938(1923) | ~ (0.6) | + (37.5) | + (27.7) | ~ (0.3) | Stockholm, Sweden | Lindsley and Grell 1968 |
| Urbana-S* | <1938 | + (2.4) | ~ (21.9) | ~ (5.5) | − (0) | Illinois, USA | Bridges†, (Lindsley and Grell, 1968) |
| Berlin-K* | <1950 | + (6.6) | + (32.1) | ~ (3.6) | − (0) | Berlin, Germany | Ruebenbauer *et al.* 2008 |
| Hikone-R* | 1950-59 | + (6.2) | ~ (17.9) | ~ (5.8) | − (0) | Japan | Galindo *et al.* 1995 |
| Florida-9* | <1952 | + (7.6) | + (31.4) | + (26.9) | + (77.3) | Florida, USA | Lindsley and Grell 1968 |
| Dmel68* | 1954 | + (14.1) | + (40.6) | + (16.2) | − (0) | Israel | NDSSC |
| B1(BER1) | 1954 | + (15.8) | + (30.0) | ~ (2.6) | − (0) | Bermuda | FlyBase |
| A3(BS1) | 1954 | + (13.7) | + (23.3) | ~ (2.4) | − (0) | Barcelona, Spain | FlyBase |
| B2(CA1) | 1954 | + (8.2) | + (33.7) | ~ (1.3) | − (0) | Capetown, South Africa | FlyBase |
| B3(QI2) | 1954 | + (10.2) | + (30.3) | + (11.1) | − (0) | Israel | FlyBase |
| A2(BOG1) | 1962 | + (18.2) | + (40.7) | + (19.5) | − (0) | Bogota, Colombia | FlyBase |
| A4(KSA2) | 1963 | + (3.0) | + (29.4) | ~ (1.3) | − (0) | Koriba Dam, Zimbabwe | FlyBase |
| B4(RVC3) | 1963 | + (15.8) | + (44.1) | + (64.7) | − (0) | California, USA | FlyBase |
| A5(VAG1) | 1965 | + (7.7) | + (31.4) | + (10.9) | − (0) | Athens, Greece | FlyBase |
| A6(wild5B) | 1966 | + (15.2) | + (31.3) | + (84.2) | − (0) | Georgia, USA | FlyBase |
| Harwich | 1967 | + (5.5) | + (55.2) | + (12.9) | + (60.1) | Massachusets, USA | NDSSC |
| Pi2* | 1975 | + (8.8) | + (31.7) | + (98.9) | + (39.7) | N.A. | Engels 1979 |
| w1118* | <1987 | + (5.2) | + (40.5) | + (35.5) | − (0) | N.A. | first used by Black *et al.* 1987 |
| AB8 (Sam;ry506) | N.A. | ~ (0.5) | ~ (28.1) | ~ (2.5) | − (0) | N.A. | N.A. |
| wk* | N.A. | + (10.4) | ~ (18.0) | ~ (4.7) | − (0) | N.A. | N.A. |
| Amherst-3* | N.A. | + (11.8) | + (35.2) | ~ (4.7) | − (0) | Massachusets, USA | N.A. |
| Iso1 | N.A. | + (20.9) | + (32.0) | + (28.6) | − (0) | N.A. | N.A. |

Table 2: Position in Tirant (pos.), reference allele (ref.) and frequency of the reference allele for SNPs with notable allele frequency differences between Iso-1 and natural populations (GDL, DrosEU and Dros-RTEC). For an overview of all SNPs in Iso-1 and some GDL lines see supplementary fig. 8.

| pos | refbase | Iso-1 (SRR1663590) | GDL-Other (SRR1663540) | GDL-Other (SRR1663560) | GDL-Other (SRR1663600) | DrosEU (SRR5647729) | DrosEU (SRR5647776) | Dros-RTEC (SRR3590550) | Dros-RTEC (SRR3939104) |
|---|---|---|---|---|---|---|---|---|---|
| 230 | C | 0.948 | 0.013 | 0 | 0 | 0.011 | 0 | 0 | 0 |
| 2485 | A | 0.792 | 0 | 0 | 0 | 0.046 | 0.036 | 0.038 | 0.057 |
| 2872 | G | 0.937 | 0 | 0 | 0 | 0.054 | 0.027 | 0.028 | 0.073 |
| 3014 | A | 0.891 | 0 | 0.167 | 0 | 0.032 | 0.023 | 0.033 | 0.05 |
| 5793 | C | 0.908 | 0 | 0 | 0 | 0 | 0 | 0 | 0 |
| 5925 | A | 0.931 | 0.134 | 0.152 | 0.227 | 0.131 | 0.14 | 0.042 | 0.103 |
| 6045 | G | 0.882 | 0 | 0 | 0 | 0 | 0.012 | 0.053 | 0.047 |
| 8337 | C | 0.931 | 0 | 0 | 0 | 0.004 | 0 | 0 | 0.003 |

Table 3: Position in Tirant (pos), reference allele (ref.) and frequency of the reference allele for SNPs with notable allele frequency differences between populations from Tasmania and other geographic locations (GDL). For an overview of all SNPs in Tasmanian and non-Tasmanian populations see supplementary fig. 9.

| pos | ref. | GDL-Tasm. (SRR1663590) | GDL-Tasm. (SRR1663591) | GDL-Tasm. (SRR1663592) | GDL-Other (SRR1663540) | GDL-Other (SRR1663560) | GDL-Other (SRR1663600) |
|---|---|---|---|---|---|---|---|
| 275 | T | 0.071 | 0.04 | 0.097 | 0.93 | 0.336 | 0.873 |
| 3921 | G | 0.075 | 0.111 | 0.084 | 0.68 | 0.824 | 0.791 |
| 5091 | T | 0.396 | 0.682 | 0.923 | 1 | 1 | 1 |
| 6757 | A | 0.932 | 1 | 0.993 | 0.734 | 0.366 | 0.439 |
| 8382 | T | 0.068 | 0.066 | 0.137 | 0.871 | 0.354 | 0.788 |

Table 4: Number of dysgenic and not-dysgenic ovaries in the F1 of reciprocal crosses between a strain having recent Tirant insertions (Urbana-S) and a strain not having recent Tirant insertions (Lausanne-S). Crosses were performed at two temperatures and three replicates were used for each cross. The direction of the cross had no significant influence on the fraction of dysgenic ovaries at both temperatures (Cochran–Mantel–Haenszel test; $p_{25} = 0.736$, $p_{29} = 0.742$).

| female | male | temp. | rep. | not-dysgenic | dysgenic |
|---|---|---|---|---|---|
| Urbana-S | Lausanne-S | 25°C | 1 | 17 | 0 |
| Urbana-S | Lausanne-S | 25°C | 2 | 13 | 0 |
| Urbana-S | Lausanne-S | 25°C | 3 | 13 | 0 |
| Urbana-S | Lausanne-S | 29°C | 1 | 12 | 1 |
| Urbana-S | Lausanne-S | 29°C | 2 | 18 | 0 |
| Urbana-S | Lausanne-S | 29°C | 3 | 18 | 0 |
| Lausanne-S | Urbana-S | 25°C | 1 | 13 | 0 |
| Lausanne-S | Urbana-S | 25°C | 2 | 15 | 0 |
| Lausanne-S | Urbana-S | 25°C | 3 | 15 | 0 |
| Lausanne-S | Urbana-S | 29°C | 1 | 18 | 0 |
| Lausanne-S | Urbana-S | 29°C | 2 | 11 | 0 |
| Lausanne-S | Urbana-S | 29°C | 3 | 15 | 0 |

Table 5: Publicly available short read data used in this work

| | |
|---|---|
| Mackay et al. (2012) | SRR018294, SRR018305, SRR018517, SRR018521, SRR018574, SRR018580, SRR018582, SRR018593, SRR018601, SRR834538, SRR834536, SRR834540, SRR834542, SRR834550, SRR834548, SRR048925, SRR834549, SRR834524, SRR834525, SRR834528, SRR834529, SRR834531, SRR834532, SRR834533, SRR834534, SRR834535, SRR834504, SRR834530, SRR834510, SRR834505, SRR834506, SRR834507, SRR834513, SRR834515, SRR834516, SRR834544, SRR834539, SRR834554, SRR834520, SRR835025, SRR835023, SRR835028, SRR835026, SRR835029, SRR051592, SRR051594, SRR835024, SRR835027, SRR835031, SRR835032, SRR835033, SRR835037, SRR835035, SRR835036, SRR835038, SRR835040, SRR835041, SRR835042, SRR835045, SRR835046, SRR051896, SRR835081, SRR835101, SRR051905, SRR835082, SRR835083, SRR835085, SRR835099, SRR835088, SRR835089, SRR835091, SRR835092, SRR835093, SRR835094, SRR835095, SRR835051, SRR835052, SRR835057, SRR835053, SRR835064, SRR835065, SRR835066, SRR835070, SRR835067, SRR835068, SRR835071, SRR835074, SRR835075, SRR835076, SRR835078, SRR835080, SRR835084, SRR835069, SRR835073, SRR834546, SRR835103, SRR060062, SRR835100, SRR060098, SRR835090, SRR835058, SRR834537, SRR834543, SRR835030, SRR835049, SRR060821, SRR835072, SRR834541, SRR835096, SRR835054, SRR834521, SRR834517, SRR834545, SRR835050, SRR933564, SRR835223, SRR835228, SRR835236, SRR835242, SRR835252, SRR835256, SRR933571, SRR933572, SRR933575, SRR933577, SRR835333, SRR933578, SRR933580, SRR933582, SRR933585, SRR933586, SRR933587, SRR933588, SRR933591, SRR933595, SRR835338, SRR933596, SRR933597, SRR933598, SRR933600, SRR835247, SRR933563, SRR932121, SRR835221, SRR933566, SRR933569, SRR933570, SRR933573, SRR835331, SRR835326, SRR933581, SRR933589, SRR933593, SRR933594, SRR835341, SRR835345, SRR933599, SRR835349, SRR835329, SRR933583, SRR933592, SRR933601, SRR933579, SRR835343, SRR835939, |
| Lack et al. (2015) | SRR189040, SRR1686796, SRR1688222, SRR189389, SRR306623, SRR306611, SRR203502, SRR306616, SRR306618 |
| Bergland et al. (2014) | SRR1525685, SRR1525694, SRR1525695, SRR1525696, SRR1525697, SRR1525698, SRR1525699, SRR1525768, SRR1525769, SRR1525770, SRR1525771, SRR1525772, SRR1525773, SRR1525774, SRR2006283 |
| Kapun et al. (2018) | SRR5647729, SRR5647730, SRR5647731, SRR5647732, SRR5647733, SRR5647734, SRR5647735, SRR5647736, SRR5647737, SRR5647738, SRR5647739, SRR5647740, SRR5647741, SRR5647742, SRR5647743, SRR5647744, SRR5647745, SRR5647746, SRR5647747, SRR5647748, SRR5647749, SRR5647750, SRR5647751, SRR5647752, SRR5647753, SRR5647754, SRR5647755, SRR5647756, SRR5647757, SRR5647758, SRR5647759, SRR5647760, SRR5647761, SRR5647762, SRR5647763, SRR5647764, SRR5647765, SRR5647766, SRR5647767, SRR5647768, SRR5647769, SRR5647770, SRR5647771, SRR5647772, SRR5647773, SRR5647774, SRR5647775, SRR5647776 |
| Machado et al. (2019) | SRR3590550, SRR3590551, SRR3590554, SRR3590555, SRR3590556, SRR3590557, SRR3590558, SRR3590559, SRR3590560, SRR3590561, SRR3590562, SRR3590563, SRR3939042, SRR3939043, SRR3939044, SRR3939045, SRR3939046, SRR3939047, SRR3939048, SRR3939049, SRR3939050, SRR3939051, SRR3939052, SRR3939054, SRR3939056, SRR3939057, SRR3939058, SRR3939059, SRR3939076, SRR3939077, SRR3939078, SRR3939080, SRR3939081, SRR3939082, SRR3939083, SRR3939084, SRR3939085, SRR3939086, SRR3939087, SRR3939088, SRR3939089, SRR3939091, SRR3939092, SRR3939093, SRR3939094, SRR3939095, SRR3939096, SRR3939097, SRR3939098, SRR3939099, SRR3939100, SRR3939101, SRR3939102, SRR3939103, SRR3939104, SRR8061818, SRR8061819 |
| Grenier et al. (2015) | SRR1663528, SRR1663529, SRR1663530, SRR1663531, SRR1663532, SRR1663533, SRR1663534, SRR1663535, SRR1663536, SRR1663537, SRR1663538, SRR1663539, SRR1663540, SRR1663541, SRR1663542, SRR1663543, SRR1663544, SRR1663545, SRR1663546, SRR1663547, SRR1663548, SRR1663549, SRR1663550, SRR1663551, SRR1663552, SRR1663553, SRR1663554, SRR1663555, SRR1663556, SRR1663557, SRR1663558, SRR1663559, SRR1663560, SRR1663561, SRR1663562, SRR1663563, SRR1663564, SRR1663565, SRR1663566, SRR1663567, SRR1663568, SRR1663569, SRR1663570, SRR1663571, SRR1663572, SRR1663573, SRR1663574, SRR1663575, SRR1663576, SRR1663577, SRR1663578, SRR1663579, SRR1663580, SRR1663581, SRR1663582, SRR1663583, SRR1663584, SRR1663585, SRR1663586, SRR1663587, SRR1663588, SRR1663589, SRR1663590, SRR1663591, SRR1663592, SRR1663593, SRR1663594, SRR1663595, SRR1663596, SRR1663597, SRR1663598, SRR1663599, SRR1663600, SRR1663601, SRR1663602, SRR1663603, SRR1663604, SRR1663605, SRR1663606, SRR1663607, SRR1663608, SRR1663609, SRR1663610, SRR1663611 |
| Wierzbicki et al. (2020) | SRR11460805, SRR11460802, SRR11460799 |
| Jakšić et al. (2017) | SRR5851905, SRR5851906 |
| Garrigan et al. (2014) | SRR1560275 |
| Turissini et al. (2015) | SRR1774232 |
| Hill et al. (2016) | SRR3113258 |
| Schrider et al. (2018) | SRR5860571, SRR5860572, SRR5860576, SRR5860577, SRR5860582 |
| Lanno et al. (2019) | SRR5860615, SRR5860617, SRR5860619, SRR5860620, SRR5860621, SRR5860622 |
| Miller et al. (2018) | SRR6425993 |
| Kang et al. (2019) | SRR6714726 |
| Melvin et al. (2018) | SRR7698174 |
| Meany et al. (2019) | SRR8834567, SRR8834568, SRR8834569 |
| Cooper et al. (2019) | SRR8840592 |
| Garrigan et al. (2012) | SRR9030358, SRR9030360 |
| Rogers et al. (2014) | SRR9699990, SRR9700000 |
| Stewart and Rogers (2019) | SRR9700024, SRR9700028, SRR9700036, SRR9700038, SRR9700041, SRR9700049, SRR9700063 |

### References


Bergland, A. O., Behrman, E. L., O'Brien, K. R., Schmidt, P. S., and Petrov, D. A. (2014). Genomic Evidence of Rapid and Stable Adaptive Oscillations over Seasonal Time Scales in *Drosophila*. *PLoS Genetics*, 10(11).

Chakraborty, M., Emerson, J. J., Macdonald, S. J., and Long, A. D. (2019). Structural variants exhibit widespread allelic heterogeneity and shape variation in complex traits. *Nature Communications*, 10(1):419275.

Cooper, J. C., Guo, P., Bladen, J., and Phadnis, N. (2019). A triple-hybrid cross reveals a new hybrid incompatibility locus between *D. melanogaster* and *D. sechellia*. *bioRxiv*, page 590588.

Drosophila 12 Genomes Consortium (2007). Evolution of genes and genomes on the *Drosophila* phylogeny. *Nature*, 450(7167):203–18.

Garrigan, D., Kingan, S. B., Geneva, A. J., Andolfatto, P., Clark, A. G., Thornton, K. R., and Presgraves, D. C. (2012). Genome sequencing reveals complex speciation in the *Drosophila simulans* clade. *Genome Research*, 22(8):1499–1511.

Garrigan, D., Kingan, S. B., Geneva, A. J., Vedanayagam, J. P., and Presgraves, D. C. (2014). Genome diversity and divergence in *Drosophila mauritiana*: Multiple signatures of faster X evolution. *Genome Biology and Evolution*, 6(9):2444–2458.

Grenier, J. K., Roman Arguello, J., Moreira, M. C., Gottipati, S., Mohammed, J., Hackett, S. R., Boughton, R., Greenberg, A. J., and Clark, A. G. (2015). Global diversity lines-a five-continent reference panel of sequenced *Drosophila melanogaster* strains. *G3: Genes, Genomes, Genetics*, 5(4):593–603.

Hill, T., Schlötterer, C., and Betancourt, A. J. (2016). Hybrid Dysgenesis in *Drosophila simulans* Associated with a Rapid Invasion of the P-Element. *PLoS Genetics*, 12(3):1–17.

Hoskins, R. A., Carlson, J. W., Wan, K. H., Park, S., Mendez, I., Galle, S. E., Booth, B. W., Pfeiffer, B. D., George, R. A., Svirskas, R., et al. (2015). The Release 6 reference sequence of the *Drosophila melanogaster* genome. *Genome research*, 25(3):445–458.

Jakšić, A. M., Kofler, R., and Schlötterer, C. (2017). Regulation of transposable elements: Interplay between TE-encoded regulatory sequences and host-specific trans-acting factors in *Drosophila melanogaster*. *Molecular Ecology*, 26(19):5149–5159.

Kang, L., Rashkovetsky, E., Michalak, K., Garner, H. R., Mahaney, J. E., Rzigalinski, B. A., Korol, A., Nevo, E., and Michalak, P. (2019). Genomic divergence and adaptive convergence in *Drosophila simulans* from Evolution Canyon, Israel. *Proceedings of the National Academy of Sciences*, 116(24):11839 – 11844.

Kapun, M., Aduriz, M. G. B., Staubach, F., Vieira, J., Obbard, D., Goubert, C., Stabelli, O. R., Kankare, M., Haudry, A., Wiberg, R. A. W., et al. (2018). Genomic analysis of European *Drosophila melanogaster* populations on a dense spatial scale reveals longitudinal population structure and continent-wide selection. *bioRxiv*, page 313759.

Lack, J. B., Cardeno, C. M., Crepeau, M. W., Taylor, W., Corbett-Detig, R. B., Stevens, K. A., Langley, C. H., and Pool, J. E. (2015). The *Drosophila* genome nexus: a population genomic resource of 623 *Drosophila melanogaster* genomes, including 197 from a single ancestral range population. *Genetics*, 199(4):1229–1241.

Lanno, S. M., Shimshak, S. J., Peyser, R. D., Linde, S. C., and Coolon, J. D. (2019). Investigating the role of Osiris genes in *Drosophila sechellia* larval resistance to a host plant toxin. *Ecology and Evolution*, 9(4):1922–1933.

Lindsley, D. H. and Grell, E. H. (1968). *Genetic variations of Drosophila melanogaster*. Carnegie Institute of Washington Publication.

Machado, H. E., Bergland, A. O., Taylor, R., Tilk, S., Behrman, E., Dyer, K., Fabian, D. K., Flatt, T., González, J., Karasov, T. L., et al. (2019). Broad geographic sampling reveals predictable, pervasive, and strong seasonal adaptation in *Drosophila*. *bioRxiv*, page 337543.

Mackay, T. F., Richards, S., Stone, E. A., Barbadilla, A., Ayroles, J. F., Zhu, D., Casillas, S., Han, Y., Magwire, M. M., Cridland, J. M., et al. (2012). The *Drosophila melanogaster* genetic reference panel. *Nature*, 482(7384):173–178.

Meany, M. K., Conner, W. R., Richter, S. V., Bailey, J. A., Turelli, M., and Cooper, B. S. (2019). Loss of cytoplasmic incompatibility and minimal fecundity effects explain relatively low Wolbachia frequencies in *Drosophila mauritiana*. *Evolution*, 73(6):1278–1295.

Melvin, R. G., Lamichane, N., Havula, E., Kokki, K., Soeder, C., Jones, C. D., and Hietakangas, V. (2018). Natural variation in sugar tolerance associates with changes in signaling and mitochondrial ribosome biogenesis. *eLife*, 7:e40841.

Miller, D. E., Staber, C., Zeitlinger, J., and Hawley, R. S. (2018). Highly Contiguous Genome Assemblies of 15 *Drosophila* Species Generated Using Nanopore Sequencing . *G3; Genes|Genomes|Genetics*, 8(10):3131–3141.

Riddle, N. C., Minoda, A., Kharchenko, P. V., Alekseyenko, A. A., Schwartz, Y. B., Tolstorukov, M. Y., Gorchakov, A. A., Jaffe, J. D., Kennedy, C., Linder-Basso, D., et al. (2011). Plasticity in patterns of histone modifications and chromosomal proteins in *Drosophila* heterochromatin. *Genome Research*, 21(2):147–163.

Rogers, R. L., Cridland, J. M., Shao, L., Hu, T. T., Andolfatto, P., and Thornton, K. R. (2014). Landscape of standing variation for tandem duplications in *Drosophila yakuba* and *Drosophila simulans*. *Molecular biology and evolution*, 31(7):1750–1766.

Schrider, D., Ayroles, J., Matute, D., and Kern, A. (2018). Supervised machine learning reveals introgressed loci in the genomes of *Drosophila simulans* and *D. sechellia*. *PLOS Genetics*, 14(4):170670.

Stewart, N. B. and Rogers, R. L. (2019). Chromosomal rearrangements as a source of new gene formation in *Drosophila yakuba*. *PLOS Genetics*, 15(9):e1008314.

Turissini, D. A., Liu, G., David, J. R., and Matute, D. R. (2015). The evolution of reproductive isolation in the *Drosophila yakuba* complex of species. *Journal of Evolutionary Biology*, 28(3):557–575.

Wierzbicki, F., Schwarz, F., Cannalonga, O., and Kofler, R. (2020). Generating high quality assemblies for genomic analysis of transposable elements. *bioRxiv*.
